# Pharming: Joint Clonal Tree Reconstruction of SNV and CNA Evolution from Single-cell DNA Sequencing of Tumors

**DOI:** 10.1101/2024.11.17.623950

**Authors:** Leah L. Weber, Anna Hart, Idoia Ochoa, Mohammed El-Kebir

**Author notes:** Correspondence: {, }.

## Abstract

Cancer arises through an evolutionary process in which somatic mutations, including single nucleotide variants (SNVs) and copy number aberrations (CNAs), drive the development of a malignant, heterogeneous tumor. Reconstructing this evolutionary history from sequencing data is critical for understanding the order in which mutations are acquired and the dynamic interplay between different types of alterations. Advances in modern whole genome single-cell sequencing now enable the accurate inference of copy number profiles in individual cells. However, the low sequencing coverage of these low pass sequencing technologies poses a challenge for reliably inferring the presence or absence of SNVs within tumor cells, limiting the ability to simultaneously study the evolutionary relationships between SNVs and CNAs.

In this work, we introduce a novel tumor phylogeny inference method, Pharming, that jointly infers the evolutionary histories of SNVs and CNAs. Our key insight is to leverage the high accuracy of copy number inference methods and the fact that SNVs co-occur in regions with CNAs in order to enable more precise tumor phylogeny reconstruction for both alteration types. We demonstrate via simulations that Pharming outperforms state-of-the-art single-modality tumor phylogeny inference methods. Additionally, we apply Pharming to a triple-negative breast cancer case, achieving high-resolution, joint reconstruction of CNA and SNV evolution, including the *de novo* detection of a clonal whole-genome duplication event. Thus, Pharming offers the potential for more comprehensive and detailed tumor phylogeny inference for high-throughput, low-coverage single-cell DNA sequencing technologies compared to existing approaches.

**Availability:** https://github.com/elkebir-group/Pharming

## 1 Introduction

Cancer arises as the result of an evolutionary process, wherein somatic mutations accumulate over time leading to the proliferation of cancer cells and the development of malignant heterogeneous tumors [1] (Fig. 1a). To gain insight into cancer and optimize treatments, researchers aim to reconstruct a tree, or tumor phylogeny, that depicts this evolutionary process from sequencing data. This has been accomplished via both bulk [2–4] and/or single-cell DNA sequencing data (scDNA-seq) [5–11]. The main advantage of single-cell DNA sequencing over bulk sequencing is the ability to observe individual cellular profiles, resulting in higher fidelity subclonal reconstruction. While scDNA-seq technologies are rapidly improving, their main drawback is that existing technologies typically favor analysis of only a single type of genetic alteration. For example, MissionBio Tapestri [12], a high-throughput, medium-to-high coverage, targeted scDNA-seq sequencing technology, is ideal for studying the evolutionary history of point mutations, or single-nucleotide variants (SNVs), covered by a panel of driver genes of cancer. However, accurately identifying larger somatic mutations involving amplifications and deletions of genomic regions from this technology, known as copy number aberrations (CNAs), remains an ongoing challenge. On the other hand, high-throughput low pass scDNA-seq technologies, such as direct library preparation (DLP+) [13] and acoustic cell tagmentation (ACT) [14], are ideally suited for inferring copy number profiles within tumors due to their uniform coverage of the genome. However, the sparsity of coverage makes it difficult to accurately infer the cellular profiles of SNVs (Fig. 1a,b).

**Figure 1:**
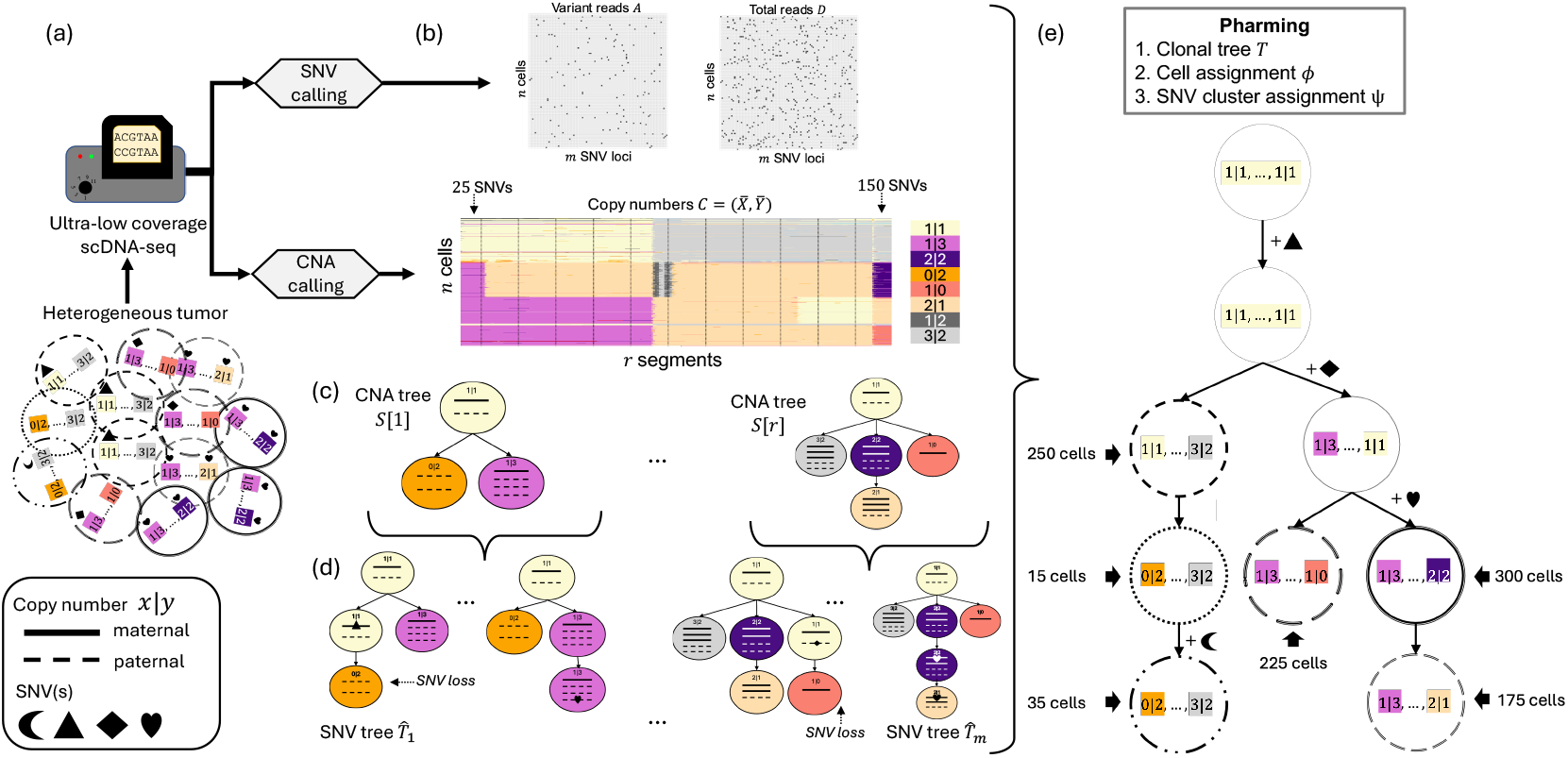
Pharming infers a clonal tree that jointly reconstructs the evolutionary history of CNAs and SNVs. (a) A heterogeneous tumor comprising both CNAs and SNVs is sequenced with low pass scDNA-seq technologies, such as DLP+ [13] and ACT [14]. (b) SNV and CNA calling produce the input to Pharming: variant and total read counts *A, D* for *n* cells and *m* SNV loci, and allele-specific copy number profiles 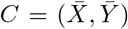 for *n* cells and *r* segments. Additionally, SNVs are mapped to the segment in which they occur. (c) The underlying evolutionary of history of CNAs with respect to the individual genomic segments 1 through *r* of the heterogeneous tumor is represented by a CNA tree *S*[*ℓ*] for segment *ℓ*. (d) The underlying evolutionary history of individual SNVs, which co-occur in the identified segments is represented by an SNV tree 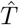,which relates the evolution of SNVs to CNAs. (e) These smaller trees are embedded within Pharming’s output clonal tree *T*, modeling the evolution of the tumor’s *m* SNVs and *r* segments. Pharming also outputs a cell assignment *ϕ*, mapping individual cells to nodes in the clonal tree, and an SNV clustering *ψ*, identifying clusters of SNVs that are introduced along the same incoming edge of the clonal tree.

Importantly, CNAs and SNVs do not evolve independently (Fig. 1c,d) and better understanding their co-evolution holds the potential for increasing our basic knowledge of cancer. There are a number of methods that have considered these dependencies. For targeted sequencing and medium-to-high coverage scDNA-seq, these methods include SCARLET [15], COnDoR [8], COMPASS [16] and SCsnvcna [17]. In SCARLET, Satas et al. [15] infer an SNV phylogeny from a CNA tree, which provides constraints on where losses can occur. However, this method requires the CNA tree as input. COnDoR [8] obviates the need for an input CNA tree by relying instead on a given clustering of cells as a proxy for copy-number profiles and proposes the constrained *k*-Dollo model that only allows SNV losses to occur when there is a shift in the cluster assignment of a node. SCsnvcna [17] is a method to place SNVs on a CNA tree inferred from cells derived from two independent sequencing experiments and is impacted by sampling biases. Additionally, SCsnvcna takes a binarized cell by SNV matrix as input and does not model read counts, which would better capture the interplay between an SNV and a CNA mutation. Lastly, COMPASS [16] infers a joint SNV and CNA phylogeny using maximum likelihood inference with Markov chain Monte Carlo (MCMC), but limits the inference of events to be either SNV, copy number gain, copy number loss or copy neutral-LOH and does not directly model the evolution of copy number profiles. Critically, since these methods use either MCMC or integer linear programming formulations, they do not scale to the number of cells and/or SNVs present in low pass scDNA-seq data. Additionally, they are unable to handle the high missing data rate resulting from the sparsity of coverage. Phertilizer [5] is a method specifically designed for low pass scDNA-seq data, exploiting the dependencies between CNAs and SNVs to guide SNV phylogeny inference via a low-dimensional embedding of the binned read counts as a surrogate for copy number signal. However, Phertilizer has two major shortcomings. First, despite loss of heterozygosity and copy number loss being ubiquitous in cancer, Phertilizer makes uses of the infinite sites assumption and therefore does not allow SNV loss due to CNAs. Second, Phertilizer does not place CNA events on its inferred phylogenies.

In this work, we overcome these limitations by developing a divide-and-conquer method called Pharming, which infers a comprehensive clonal tree with both CNA and SNV events from low-pass single-cell DNA sequencing data (Fig.1e). In the divide step, Pharming infers a set of candidate trees for each specific genomic segment or region, focusing on the limited copy number states within that segment and the SNVs that occur within that region. To accurately integrate candidate clonal trees in the merge step, Pharming uses the fact that the same set of cells is used in each segment-specific clonal tree as well as the fact that SNV gains can be grouped in a small number of clusters on the final tree. Using simulated tumors, we show Pharming faithfully reconstructs the evolutionary history of CNAs and SNVs with respect to a known ground truth, as well as placement of cells onto the clonal tree, even with coverages as low as 0.01×. On experimental data of breast [14] and ovarian [13] cancers, we showed that Pharming provides accurate and high-resolution joint reconstruction of CNA and SNV evolution, matching orthogonal validation data.

## 2 Materials and Methods

Suppose we have sequenced *n* cells and identified *m* SNVs from low pass scDNA-seq data. From this, we obtain the variant *A* = [*a*_*i,j*_] and total *D* = [*d*_*i,j*_] read counts for each cell *i* ∈ [*n*] = {1, …, *n*} and SNV *j* ∈ [*m*]. Using existing allele or haplotype-specific copy number callers, such as CHISEL [18], SIGNALS [19] or CNRein [20], we obtain copy number profiles 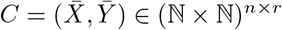 with respect to the two germline haplotypes, where *r* is the number of segments inferred during the genome segmentation by the copy number caller. More specifically, segmentation ensures the copy number call 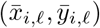 for an individual cell *i* is constant throughout all loci contained within segment *ℓ* and consequently segments may have varying size. We define an SNV-to-segment mapping *σ* : [*m*] → [*r*] that indicates the segment *ℓ* = *σ*(*j*) where SNV *j* occurs. Similarly, the set *σ*^−1^(*ℓ*) ⊆ [*m*] contains the SNVs located in segment *ℓ*. We use the tuple (*C, A, D*) as a shorthand to refer to the observed copy number profiles *C* as well as the variant read counts *A* and total read counts *D*. As SNVs can be affected by CNAs, we model the *genotype* of an SNV in an individual cell by the tuple (*x, y, w, z*), where (*x, y*) are the allele-specific copy numbers and *w* and *z* are the number of mutated copies on haplotypes *x* and *y*, respectively. We summarize key notation in Table S1.

The joint evolution of SNVs and CNAs is represented by a clonal tree. While parallel gains of an SNV are rare [21], SNV loss due to CNAs or loss-of-heterozygosity (LOH) events are common. Therefore, we make use of the Dollo evolutionary model [22] for SNVs, which allows each SNV to be gained exactly once but may be lost due to copy number loss. For CNAs, we utilize the infinite alleles assumption [23] such that each copy number state within a segment is introduced exactly once in the clonal tree. While violations of this assumption do occur in practice, it reasonably holds for larger segments and makes the challenging problem of joint clonal tree inference more tractable. This leads to the following definition of a clonal tree, illustrated in Fig. 1a and Fig. 2a.

**Figure 2:**
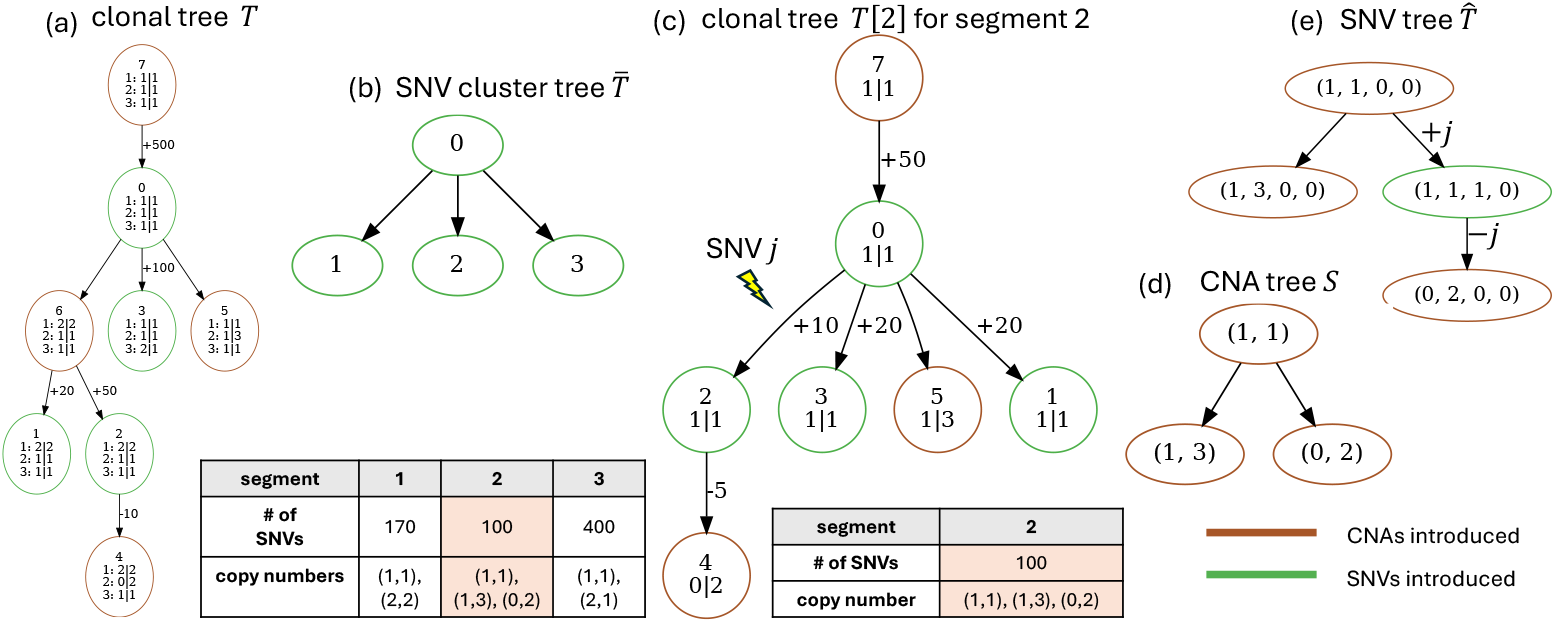
Depiction of tree structures for Pharming. (a) Pharming clonal tree *T* for *r* = 3 segments with varying number of SNVs and copy number states per segment. Newly introduced or lost SNVs are shown on the incoming edges. The copy number state for each node and each segment is labeled as “segment: *x*| *y*”. The root (7), is the normal clone. (b) Clonal tree *T* with *k* = 4 SNV clusters is consistent with SNV cluster tree 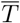. (c) Clonal tree *T* is consistent with clonal tree *T* [2] for segment 2. (d) The underlying CNA tree *S* for clonal tree *T* [2]. Clonal tree [2] is consistent with CNA tree *S*. (e) The underlying SNV tree 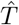 for SNV *j*, with SNV cluster assignment *ψ*(*j*) = 2.SNV tree 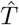 is consistent with CNA tree *S*.

### Definition 1.

A rooted tree *T* whose nodes *V* (*T*) are labeled by genotypes *g* = (*x, y, w, z*) for *r* segments, consisting of copy numbers *x* : *V* (*T*) × [*r*]→ ℕ and *y* : *V* (*T*) × [*r*] → ℕ and mutation numbers *w* : *V* (*T*) × [*m*] → ℕ and *z* : *V* (*T*) × [*m*] → ℕ, is a *clonal tree* provided (i) for each SNV *j*, there is exactly one edge (*u, v*) ∈ *E*(*T*) where the SNV was gained, i.e., where either *z*(*u, j*) = 0 and *z*(*v, j*) *>* 1, or *w*(*u, j*) = 0 and *w*(*v, j*) *>* 1; (ii) for each SNV *j* and each node *v*, the mutation number is bounded by the corresponding copy number, i.e., 0 ≤ *w*(*v, j*) ≤ *x*(*v, ℓ*) and 0 ≤ *z*(*v, j*) ≤ *y*(*v, ℓ*), where *ℓ* = *σ*(*j*) is the segment where SNV *j* is located; (iii) for each SNV *j* and each copy number 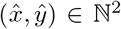 ∈ ℕ^2^, there is at most one edge (*u, v*) ∈ *E*(*T*) introducing copy number 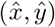 of the segment *ℓ* = *σ*(*j*), i.e., where 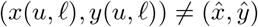 and 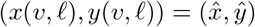.

For brevity, we use the shorthand *g*(*v, j*) = (*x, y, w, z*) to indicate (*x*(*v, σ*(*j*)), *y*(*v, σ*(*j*)), *w*(*v, j*), *z*(*v, j*)) for a node *v* and SNV *j*. The variant allele frequency (VAF) of an SNV *j*, i.e., the ratio of mutated to total copies, is particularly informative for modeling observed variant read counts conditioned on its genotype *g*(*v, j*) at node *v*.

### Definition 2.

Given a clonal tree *T* with genotypes *g*(*v, j*) = (*x, y, w, z*) the *latent variant allele frequency (VAF)* of an SNV *j* in segment *ℓ* = *σ*(*j*) of node *v* equals *f* (*v, j*) = (*w* + *z*)*/*(*x* + *y*).

We also define a cell assignment *ϕ* : [*n*] →*V* (*T*) that specifies the assignment of each cell *i* to a node *v* in clonal tree *T*. To assess goodness of fit of a clonal tree *T* and cell assignment *ϕ* with respect to the observed data (*C, A, D*), we derive a cost function, comprised of two parts.

First, we define the likelihood for the observed variant read counts *A* given total read counts *D*, clonal tree *T* and cell assignment *ϕ* under a binomial model that accounts for the probability *ϵ* of misreading a base during sequencing as 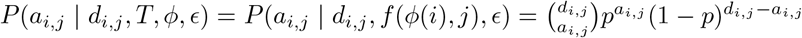, where, as derived in [5], probability 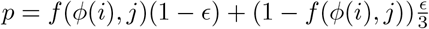 is a function of the latent VAF *f* (*ϕ*(*i*), *j*)) of SNV *j* in cell *i* and *ϵ*.

Second, while copy number calling methods are rapidly improving, they are not error-free. As a result, we permit some deviation between the observed 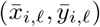 and inferred (*x*(*ϕ*(*i*), *ℓ*), *y*(*ϕ*(*i*), *ℓ*)) copy numbers. We consider a penalty term 𝒳(*i,l*) that accounts for deviations for each cell *i* and segment 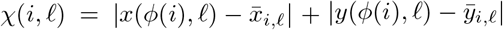. The level of tolerance for deviation is controlled by parameter *λ* ∈ℝ_≥0_. Assuming independence between segments and SNVs, we arrive at the following cost function for a given clonal tree *T* and cell assignment *ϕ*: *J*(*T, ϕ*) = − п _*i*∈[*n*]_ п _*j*∈[*m*]_ *P* (*a*_*i,j*_|*d*_*i,j*_, *T, ϕ, ϵ*) + *λ* ∑_*i*∈[*n*]_ ∑_*ℓ*∈[*r*]_ *χ*(*i, ℓ*). This leads to the following problem.

### Problem 1

(Joint Clonal Tree Inference (JCTI)). Given copy numbers 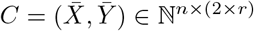,variant read counts *A* ∈ ℕ^*n*×*m*^ and total read counts *D* ∈ ℕ^*n*×*m*^ for *n* cells, *r* segments and *m* SNVs as well as parameters *λ* ∈ ℝ_≥0_ and *ϵ* ∈ [0, 1], find a clonal tree *T* labeled by genotypes *g* and cell assignment *ϕ* with minimum cost *J*(*T, ϕ*).

### 2.1 The PHARMING algorithm

One can show the JCTI problem to be NP-hard by reduction from the (Perfect Phylogeny) Flipping Problem [24]. Therefore, we develop a heuristic algorithm named Pharming that relies on the following four key insights. Firstly, similar to Phertilizer [5], we expect gains of SNVs to cluster into a small number *k* of clusters where *k* ≪ *m*. As such, we define *ψ* : [*m*] → [*k*] to be a mapping of an SNV to one of *k* SNV clusters.

Secondly, SNVs in the same SNV cluster must share a descendant cell fraction (DCF) [15]. The DCF of an SNV is defined as the fraction of cells that are assigned to or are descendants of the node where an SNV was introduced. By construction, the DCFs account for cells assigned to nodes even where an SNV may have been lost. We define a mapping δ : [*k*] → [0, 1] that yields the DCF δ(*q*) of an SNV cluster *q*. Although DCFs were originally intended to facilitate SNV clustering for bulk data, they can be inferred from a pseudobulk sample. Critically, the DCFs δ constrain the evolutionary relationships between SNV clusters and provide a means to significantly reduce the underlying space of clonal trees [2].

Thirdly, the individual problems of inferring a clonal tree individually for CNA or SNVs are both challenging. However, SNVs commonly co-occur in segments impacted by CNAs, yielding a valuable signal for the evolutionary relationships between these events. Thus, considering the interplay of the mutation types allows us to capitalize on additional signal that is unavailable when looking only on a single mutation type.

Intuitively, if we look at the evolutionary history of a single SNV, it is by Def. 1 a clonal tree. Likewise, if we look at the evolutionary history of a single segment containing a subset of SNVs that occur within that segment, it is also a clonal tree. Even further, if we *merge* two clonal trees constructed from disjoint segments containing disjoint subsets of SNVs such that the evolutionary history of each individual SNV is *consistent with* the clonal tree of origin, then the result is a clonal tree. Therefore, our fourth key insight is to capitalize on this substructure by utilizing a divide-and-conquer approach. In other words, we first focus on simpler subproblems, e.g., infer the clonal tree for a single segment, and then we *merge* the solutions of these subproblems such that *consistency* with the original solutions is mantained. Furthermore, to ensure that in the merging steps a total of *k* SNV clusters are inferred, we require clonal trees for each segment to be consistent with a fixed SNV cluster tree.

P<sc>harming</sc> utilizes these key ideas to solve the JCTI problem in a three-phased algorithm: (i) initialization (Sec. 2.1.2), (ii) clonal tree inference for a single segment (Sec. 2.1.3) and (iii) integration of clonal trees (Sec. 2.1.4). In the following, we define some additional terminology and notation helpful for describing the main phases of Pharming. While the provided description assumes a fixed number *k* of SNV clusters, a sweep across reasonable values of this parameter in conjunction with model selection is used to find a heuristic solution to the JCTI problem (Sec. A.2.1).

#### 2.1.1 Preliminaries

Since Pharming is a divide-and-conquer algorithm, it makes use of a variety of tree structures (Fig. 2), some of which are special types of clonal trees, that efficiently capture the complex evolutionary history of both SNVs and CNAs in a heterogeneous tumor. We start by formally defining these structures and the terminology that defines their relationships to one another in the inferred clonal tree *T*. Specifically, we define the key building blocks of a clonal tree: (i) a CNA tree *S*, (ii) an SNV tree 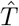 and a (iii) SNV cluster tree 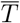.

First, a CNA tree *S* models the evolution of the copy number states of a single segment, respecting the same evolutionary constraints as a clonal tree.

##### Definition 3.

A rooted tree *S* labeled by copy numbers *x, y* : *V* (*S*) → ℕ is a *CNA tree* provided for each copy number state 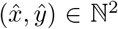 there exists at most one edge (*u, v*) ∈ *E*(*S*) where 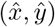 is introduced, i.e., 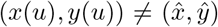 and 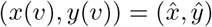.

Second, an SNV tree 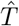 describes the evolutionary history of an individual SNV, respecting the same evolutionary constraints as a clonal tree.

##### Definition 4.

A rooted tree 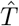 labeled by genotypes 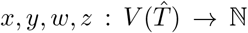 is an *SNV tree* provided (i) the SNV was gained once; (ii) the mutation number is bounded by the corresponding copy number; and (iii) each copy number 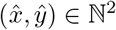 is attained at most once.

Third, we introduce an important tree structure for our algorithm referred to as an SNV cluster tree for a fixed number *k* of SNV clusters. This tree reflects the evolutionary relationships of the *k* SNV clusters (Fig. 2b).

##### Definition 5.

A rooted tree 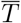 whose *k* nodes are labeled by SNV clusters is an *SNV cluster tree*.

As we show next, these three types of trees can be extracted from a clonal tree *T* on all *r* segments and *m* SNVs clustered into *k* groups. First, to extract an SNV tree 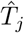 corresponding to SNV *j* on segment *ℓ* = *σ*(*j*) from *T*, we contract all edges (*u, v*) of *T* with identical genotypes for SNV locus *j*, i.e., *g*(*u, j*) = *g*(*v, j*), retaining only labels corresponding to SNV *j* and segment *ℓ* (Fig. 2e). We say that an SNV *j* of a clonal tree *T* is *consistent* with an SNV tree 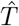 if the extracted SNV tree 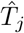 from *T* is identical to 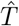.Second, we extract a CNA tree *S*[*ℓ*] from *T* corresponding to segment *ℓ* by contracting all edges (*u, v*) of *T* with identical copy numbers for segment *ℓ*, i.e., *x*(*u, ℓ*) = *x*(*v, ℓ*) and *y*(*u, ℓ*) = *y*(*v, ℓ*), retaining only labels for segment *ℓ* (Fig. 2d). We say that segment *ℓ* of a clonal tree *T* is *consistent with* with a CNA tree *S* if the extracted CNA tree *S*[*ℓ*] from *T* is identical to *S*. Moreover, we say that an SNV tree 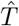 is *consistent* with a CNA tree *S* if the extracted CNA tree from 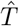 is identical to *S*. Third, we extract an SNV cluster tree 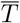 from *T* by contracting all edges where no SNVs were introduced — such that for each of the remaining *k ™*1 edges (*u, v*) there exists an SNV *j* where max {*w*(*u, j*), *z*(*u, j)*} = 0 and max {*w*(*v, j*), *z*(*v, j*)} *>* 0 (Fig. 2b). We say that a clonal tree *T* is *consistent* with an SNV cluster tree 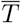 if the extracted SNV cluster tree from *T* is identical to 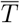. In a similar vein, we can extract a partial clonal tree *T* [*ℓ*] from *T* for each individual segment *ℓ* ∈ [*r*], or a partial clonal tree *T* [*L*] for a subset *L* ⊆ [*r*] of segments (Fig. 2c). A clonal tree *T* is *consistent* with a partial clonal tree *T* [*L*] of segments *L* ⊆ [*r*] if the extracted clonal tree from *T* with respect to segments *L* is identical to *T* [*L*].

#### 2.1.2 Initialization

We begin by initializing the DCFs δ for a fixed value of *k*. These DCFs may be obtained via an existing algorithm called DeCiFer [25] on a pseudobulk sample generated from merging the scDNA-seq data. Briefly, starting from a random initialization of *k* DCFs values, DeCiFer uses a coordinate ascent approach to alternate between (i) optimizing an assignment of each SNV *j* to a cluster *q* and an SNV tree 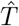 given the DCF cluster centers δ and (ii) optimizing the DCF cluster centers δ given a cluster assignment *q* and SNV tree 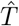 for each SNV *j*. Pharming includes a customized implementation of DeCiFer that additionally enforces consistency among underlying CNA trees for a segment (detailed in Appendix A). Given the inferred DCFs δ_1_, …, δ_*k*_ for *k* clusters, we next enumerate a set 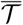 of *valid* SNV cluster trees, with the formal definition below. Additional details on enumeration of the set 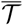 of valid SNV cluster trees are available in El-Kebir *et al*. [2].

##### Definition 6.

An SNV cluster tree 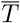 is *valid* for DCFs 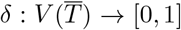 provided 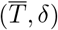 satisfy the Sum Condition(SC) [2], i.e., δ(*u*) ≥ ∑_*v*∈children(*u*)_ δ(*v*), where children(*u*) is the set of children of node *u* in 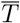.

#### 2.1.3 Clonal tree inference for a single segment

Starting from the set 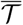 of SNV cluster trees inferred from the initial DCFs δ_1_, …, δ_*k*_, we next aim to infer a set 𝒯 [*ℓ*] of candidate clonal trees for each segment *ℓ*. Not only is each candidate clonal tree *T* [*ℓ*] ∈ 𝒯 [*ℓ*] consistent with a single SNV cluster tree 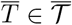, it is also consistent with a single CNA tree *S* ∈ 𝒮 [*ℓ*] — where 𝒮 [*ℓ*] is the set of CNA trees comprising the observed copy numbers of segment *ℓ* and respecting the infinite alleles assumption (see Defs. 1 and 3). Therefore, we can construct 𝒯 [*ℓ*] in a bottom-up fashion, fixing an SNV cluster tree 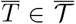 and a CNA tree *S* ∈ 𝒮 [*ℓ*] This amounts to the following subproblem, which is a variant case of the JCTI problem restricted to only *r* = 1 segment and the SNVs *σ*^−1^(*ℓ*) occurring within that single segment *ℓ* and subject to additional clonal tree consistency constraints.

##### Problem 2

(Segment Tree Inference (STI)). Given a SNV cluster tree 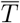 with 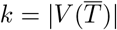 clusters, CNA tree *S* and a fixed segment *ℓ* with *m*_*ℓ*_ = |*σ*^−1^(*ℓ*)| SNVs, associated copy numbers 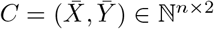,variant read counts 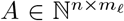 and total read counts 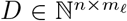, enumerate all clonal trees 𝒯 [*ℓ*] that are consistent with 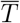 and *S* and identify genotypes *g* and cell assignments *ϕ* with minimum cost *J*(*T* [*ℓ*], *ϕ*) for each *T* [*ℓ*] ∈ 𝒯 [*ℓ*].

To solve the STI problem, we first enumerate a set 𝒯 [*ℓ*] of clonal trees consistent with both the fixed CNA tree *S* and the fixed SNV cluster tree 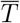, but not containing any individual SNVs nor their genotypes (Fig. S1). As each such clonal tree *T* [*ℓ*] ∈ 𝒯 [*ℓ*] only indicates where each SNV cluster *q* is introduced with respect to the copy number states in CNA tree *S*, it does not yet define the genotypes *g* of the *m*_*ℓ*_ SNVs. We must therefore find the optimal genotypes *g* and cell assignment *ϕ* for each *T* [*ℓ*] ∈ 𝒯 [*ℓ*]. We accomplish this using a coordinate descent algorithm where we alternately optimize the genotypes *g* and the cell assignment *ϕ*.

We start the coordinate descent algorithm by initializing the genotypes *g*. Given a CNA tree *S*, we enumerate a set 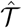 of SNV trees that are consistent with *S*. Note that the SNV assignment *ψ*(*j*) = *q* and the tree *T* [*ℓ*] constrain where the gain of an SNV occurred relative to prior CNA states. Therefore, allowed SNV trees 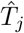 are constrained to a subset 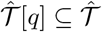 of the enumerated SNV trees. Given SNV cluster *ψ*(*j*) and SNV tree 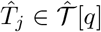,then the genotype *g*(*u, j*) of SNV *j* for every node *u ∈ V* (*T* [*ℓ*]) is uniquely determined. In Appendix A, we describe our procedure to initialize genotypes *g* by assigning each SNV to a cluster *q* and an SNV tree 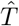 that is consistent with clonal tree *T* [*ℓ*] and SNV cluster *ψ*(*j*).

Next, we optimize the cell assignment *ϕ* given clonal tree *T* [*ℓ*] labeled by genotypes *g*. This is accomplished by computing the cost of each possible cell assignment *ϕ*(*i*) = *u* for each node *u* ∈ *V* (*T* [*ℓ*]) and then assigning cell *i* to the node with minimum cost. Similarly, given a fixed cell assignment *ϕ*, we optimize the genotypes *g* by computing the cost of each SNV cluster assignment *ψ*(*j*) = *q* and SNV tree assignment 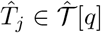 for each SNV cluster *q* ∈ [*k*] and updating the genotypes *g*(*u, j*) for each node *u* ∈*T* [*ℓ*]. We repeat this alternating optimization until cost *J*(*T* [*ℓ*], *ϕ*) converges or we reach a maximum number of iterations.

In summary, we solve the STI problem for each combination 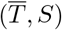 of SNV cluster trees 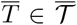 and CNA tree *S* ∈ 𝒮 [*ℓ*], retaining all enumerated pairs (*T* [*ℓ*], *ϕ*) of clonal trees and cell assignments ordered by their costs *J*(*T* [*ℓ*], *ϕ*).

#### 2.1.4 Integration of clonal trees

Given two clonal trees *T*_1_ and *T*_2_ constructed from the same underlying SNV cluster tree 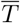 on disjoint sets *L*_1_ and *L*_2_ of segments, we consider the problem of how to ‘merge’ these trees together in order to infer a clonal tree *T* [*L*] for segments *L* = *L*_1_ ∪ *L*_2_.

**Problem 3** (Tree Merging Inference (TMI)). Given two clonal trees *T*_1_ and *T*_2_ constructed from the same underlying SNV cluster tree 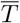 on disjoint sets *L*_1_ and *L*_2_ of segments, respectively, and *L* = *L*_1_ ∪*L*_2_ of segments, with *m*_*L*_ = |_*ℓ*∈*L*_ *σ*^−1^(*ℓ*) | SNVs, associated copy numbers 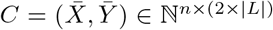,variant read counts 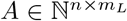 and total read counts 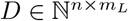, find a clonal tree *T* [*L*] labeled by genotypes *g* and cell assignment *ϕ* with minimum cost *J*(*T* [*L*], *ϕ*) such that *T* [*L*] is consistent with clonal tree *T*_1_, clonal tree *T*_2_ and SNV cluster tree 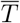.

To solve the TMI problem, we start by enumerating a set 𝒯 [*L*] of clonal trees such that each clonal tree *T* ∈ [*L*] 𝒯 [*L*] is consistent with each CNA tree *S*[*ℓ*] of each segment *ℓ ∈ L*_1_ ∪*L*_2_ and consistent with SNV cluster tree 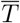. Although this sounds complicated, the key idea to solving the TMI problem is that by construction both clonal trees are consistent with the same underlying SNV cluster tree 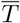. Consequently, we only need to resolve the evolutionary relationships of copy number states across distinct segments with respect to the SNV clusters. For example, if copy number 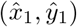 in segment *ℓ* and copy number 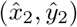 in segment *ℓ*^*′*^ are both introduced after SNV cluster *q* but before SNV *q*^*′*^, then the set 𝒯 [*L*] will include a clonal tree with copy number 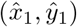 introduced prior to 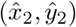 and a clonal tree with copy number 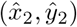 introduced before 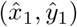. For each clonal tree *T* [*L*] ∈ 𝒯 [*L*], we find cell assignment and the *J*(*T* [*L*], *ϕ*) by assigning each cell *i* to the node *u*∈ *V* (*T* [*L*]) that minimizes costs.

To integrate a set {*T* [1], …, *T* [*r*]} of clonal trees with the same underlying SNV cluster tree 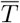, we progressively solve the TMI problem. As there are *r*! orderings, we use a heuristic where we randomly sample the *r* segments without replacement, weighted by the number *m*_*ℓ*_ = |*σ*^−1^(*ℓ*) | of SNVs in segment *ℓ*. Progressive merging may be repeated for multiple random permutations. While we leave the study of the optimal ordering strategy for future work, we seek to overcome suboptimal orderings by retaining the top *n* solutions to the TMI problem at each stage. This allows us to merge the cross product of the 𝒯 [*L*] ×𝒯 [*L*^*′*^], retaining the top *n* clonal trees for subsequent merging.

### 2.2 Availability

As the number *k* of SNV clusters is unknown, Pharming is run for reasonable values of *k* and we use the Bayesian information criterion, i.e., BIC(*k*), in a model selection stage (Appendix A.2.1). Pharming outputs the clonal tree *T* and cell assignment *ϕ* for the number *k*^∗^ = arg min_*k ∈K*_ BIC(*k*) of SNV clusters. Pharming is implemented in Python and C++, is open source, and available at https://github.com/elkebir-group/Pharming.

## 3 Results

### 3.1 In silico experiments

To evaluate Pharming, we generated *in silico* experiments with known ground truth clonal trees and cell clusterings. We set simulation parameters to *n* = 1000 cells, *m* = 5000 SNVs, *r* = 25 segments, *k* = 5 mutation clusters, coverage varying in {0.01, 0.05, 0.25} and the copy number error rate in {0, 0.035} with 5 replicates for each coverage, yielding a total of 30 experiments. For all experiments, the per base sequencing error rate *ϵ* was set to 0.001. We ran Pharming for different values of *k* ∈ {3, …, 6} SNV clusters. During the integration stage, we maintained a set of the top 5 solutions and utilized only one random ordering for progressive integration. We utilized a penalty of *λ* = 1000 in order to compute our cost function and then used BIC in order to select an inferred clonal tree *T*^∗^ and cell assignment *ϕ*^∗^ across varying number *k* ∈ {3, …, 6} of SNV clusters.

Although we note that no other existing methods infer a joint CNA and SNV clonal tree for low pass scDNA-seq data, we benchmarked Pharming against three methods that solve related problems, applying adaptations to these approaches where necessary to obtain similar outputs for comparison. First, while Phertilizer utilizes CNA signal, it infers only an SNV clonal tree under the ISA, not supporting loss and not accounting for SNV multiplicity beyond presence/absence. Second, we benchmarked against a commonly used *ad hoc* method [13], which we previously referred to as the Baseline+SCITE method [5]. Briefly, Baseline first finds a cell clustering (via UMAP and density-based clustering) and then obtains SNV genotypes from pseudobulk samples composed of cell clusters. These observed genotypes are then provided to SCITE [7] to yield an SNV clonal tree subject to the same limitations as Phertilizer. Third, we compared against Lazac [26], which infers copy number phylogenies (where each cell is a separate leaf) using the zero-agnostic copy number transformation distance. Importantly, as the first two methods do not infer the copy number states of ancestral clones, we obtain consensus copy number profiles of the inferred cell clusters. Since Lazac does not infer SNV genotypes and does not cluster cells into clones, we only include Lazac in comparisons focusing on the inference of CNA evolution. To assess performance with respect to inference of SNVs, CNAs and cell assignment, we utilized a number of performance metrics. We focus our discussion on simulations with a copy number error rate of 0.035 as these instances are more reflective of real data. Further details on simulation setup, benchmarking methods and performance metrics are provided in Appendix B.1 and illustrated in Figs. S2–S5.

To simultaneously assess the accuracy of the cell clustering and inferred evolutionary relationships between clones, we compute the *cell placement accuracy*, which is the weighted average of the ancestral pair recall, incomparable pair recall, and clustered pair recall metrics [2] for pairs of cells. A *cell placement accuracy* of 1 implies that the inferred cell placement perfectly matches ground truth. As shown in Fig. 3a, Pharming (median: 0.90) significantly outperformed both Phertilizer (median: 0.58) and Baseline+SCITE (median: 0.70). Upon further examination, we found that Phertilizer tended to underestimate the number of cell clusters (median inferred: 5 vs. median ground truth 9), resulting in a less refined cell clustering. We also observed that while the Baseline+SCITE method accurately clustered the cells based on the simulated copy number profiles (ARI: median: 0.98 vs Pharming median: 0.97), it ultimately performed poorly in correctly inferring the evolutionary relationships between clones (Figs. S6, S7). In particular, Baseline+SCITE performs especially poorly on ancestral pair recall (median: 0.03) when compared to Pharming (median: 0.96).

**Figure 3:**
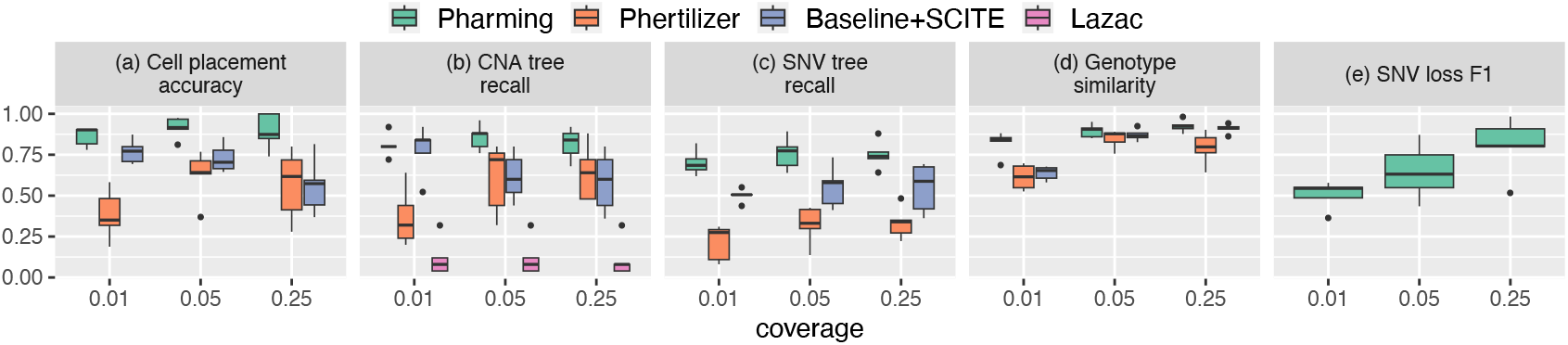
Pharming outperforms existing methods on *in silico* experiments. Benchmarking comparison of performance with known ground truth. Results are shown for *in silico* experiments with 5000 SNVs, 1000 cells, CNA error rate 0.035,and coverage varying in {0.01×, 0.05×, 0.25×} on the following metrics (a) cell placement accuracy, CNA tree recall, (c) SNV tree recall, (d) genotype similarity and (e) SNV loss F1 (defined in Appendix B.2).

To further investigate performance of clonal tree inference, we assessed the accuracy of the evolution of copy number states. We define the *CNA tree recall* as the proportion of segments where the inferred segmental CNA tree exactly matches the ground truth segmental CNA tree. Overall, Pharming (median:0.8) performed best on this metric, maintaining similar performance across the varying coverage (Fig. 3b). Interestingly, while Baseline+SCITE has a high median *CNA tree recall* of 0.72, there is a noticeable drop off in performance as sequencing coverage increases. We also observed this same phenomenon for *cell placement accuracy*. This suggests that potential false positives resulting from the pseudobulk approach to SNV genotyping may yield misleading mutation trees that do not accurate reflect the evolution of the tumor clones. In contrast, Phertilizer (median: 0.48) and Lazac (median: 0.08) had the lowest overall performance on this metric. While Phertilizer uses the cell embedding based on copy number profiles to aid in SNV tree inference, it does not directly consider the joint evolution of CNAs and SNVs. On the other hand, Lazac does not make use of any SNV information. Thus, by not performing joint inference, both methods struggle with properly reconstructing the evolution of CNA states.

We next assessed the accuracy of the joint inference of the CNAs and SNVs. Like CNA tree recall, we define *SNV tree recall* as the proportion of SNVs whose inferred SNV tree matches the ground-truth SNV tree (both in terms of topology and genotypes, modulo SNV multiplicity and allele specificity). Pharming had the highest median SNV tree recall (0.73), followed by Baseline+SCITE (median: 0.51) then Phertilizer (median: 0.3) — see Fig. 3c. Importantly, for Pharming we did not observe a significant drop off in performance as coverage decreases, highlighting the utility of our method in the ultra-low coverage regime.

Next, to simultaneously evaluate the accuracy of clonal genotyping and cell clustering, we define *genotype similarity* as 1 minus the normalized Hamming distance between the ground truth and inferred cellular genotypes when ignoring allele-specificity of the SNV. As shown in Fig. 3d, Pharming had the highest overall *genotype similarity* with a median of 0.88, closely followed by Baseline+SCITE (median: 0.86) and then by Phertilizer (median: 0.76). While Pharming and Baseline+SCITE had similar performance at coverages of 0.05× and 0.25×, we observe a significant difference in median *genotype similarity* at the lowest coverage of 0.01× (Pharming: 0.84 vs Baseline+SCITE: 0.65).

Finally, since Pharming does not assume the infinite sites model, we evaluated its ability to accurately infer SNV loss. We used a metric called *SNV loss F1*, which is the harmonic mean of two components: *SNV loss recall*–—the proportion of SNVs that undergo loss in the ground truth clonal tree and are correctly identified as undergoing loss in the inferred tree — and *SNV loss precision*— the proportion of inferred SNVs undergoing loss that actually underwent loss in the ground truth tree. Pharming had an overall median *SNV loss F1* of 0.57 (Fig. 3e), although performance increases as coverage increases (0.01 ×: 0.54, 0.05× : 0.63, 0.25 × : 0.80). Additionally, we observed that overall *SNV loss recall* was high (median: 0.85) while *SNV loss precision* was low (median: 0.57). Thus, while there is still opportunity for improving inference of SNV loss, Pharming is able to correctly infer the loss of a high proportion of SNVs.

In summary, these performance metrics highlight the capability of Pharming to more accurately infer a joint CNA and SNV clonal tree than competing methods, even at a coverage as low as 0.01 ×. We observe similar trends and performance for instances with a copy number error rate of 0 (Fig. S11).

### 3.2 Triple negative breast cancer tumors

To demonstrate the capabilities of Pharming on tumors sequenced with low pass scDNA-seq, we analyzed two triple negative breast cancer tumors, TN1 and TN3, sequenced with acoustic cell tagmentation (ACT) [14], with *n* = 1100 and *n* = 1101 cells, respectively. Both tumors had extremely low sequencing coverage with TN1 having 0.031 ×while TN3 had 0.02 ×coverage. To obtain variant and total read counts *A, D*, we called SNVs on a pseudobulk sample of each tumor, resulting in *m* = 12,118 and *m* = 12,662 SNVs for TN1 and TN3, respectively. We then ran CNRein [20] to obtain the segmented haplotype-specific copy number calls 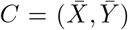. TN1 had *r* = 335 segments while TN3 had lower copy number heterogeneity resulting in only *r* = 171 segments. Additional details on these preprocessing steps are provided in Appendix B.3.1.

For each number *k* ∈ {2, 3, 4} of SNV clusters, we ran Pharming on the TN3 dataset. Using BIC analysis, we selected the clonal tree *T*^∗^ corresponding to *k* = 3 SNV clusters (Fig. 4a). Of the *m* = 12,662 SNVs, 11,798 (93%) were placed in the trunk of the clonal tree. The remaining 864 SNVs formed a single subclonal cluster. Compared to Baseline, Pharming was able to tease additional clonal structure for this tumor (Fig. S13). We assessed goodness of fit of clonal tree *T*^∗^ via the previously introduced *cell mutational burden* (CMB) metric [5] — see Appendix B.2.2 for more details on this metric. Briefly, CMB(*i, M*) is the fraction of mapped SNV loci *M* with mapped variant reads in cell *i*. Thus, we expect cells mapped to nodes *within* the clade where the set *M* of SNVs were introduced to have a CMB greater than 0, while the CMB for cells outside of the clade should be close to 0. The subclonal SNV cluster with |*M*| = 864 SNVs is introduced at clade A in clonal tree *T*^∗^ and indeed, we observed a high median within-clade CMB of 0.29 while the median CMB outside of the clade was 0, providing supporting evidence for this subclonal SNV cluster (Fig. 4b). However, the lack of other subclonal SNV clusters made this dataset less than ideal to showcase Pharming’s ability to resolve the ordering of subclonal CNAs as it is best suited for datasets with both subclonal SNV and CNA structure.

**Figure 4:**
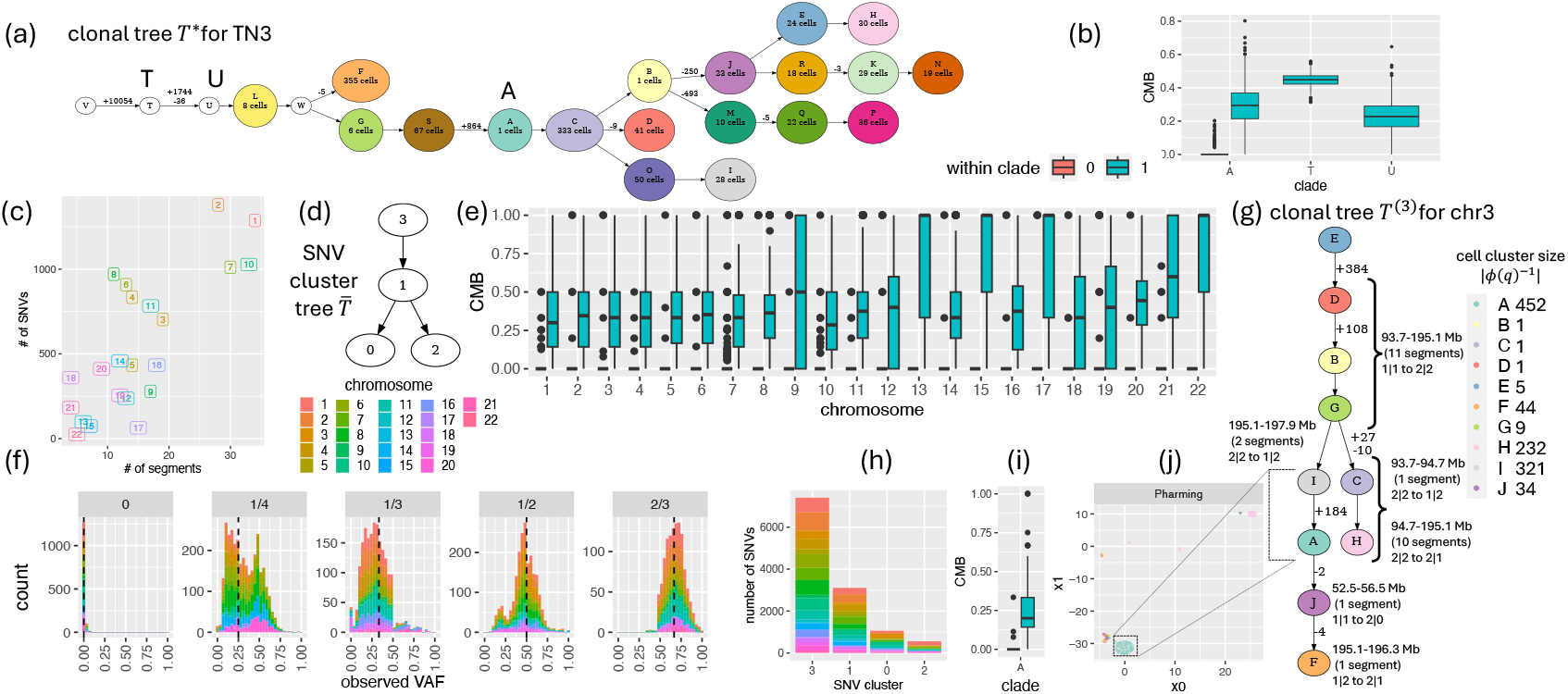
Pharming infers clonal trees for triple negative breast cancer tumors. For TN3 data: (a) Inferred clonal tree. (b) Distribution of the cell mutational burden (CMB) by clade for the TN3 clonal tree. For TN1 data: (c) Relationship between the number of identified SNVs and inferred segments by chromosome. (d) SNV cluster tree inferred by Pharming. (e) Distribution of CMB aggregated over the clades in the inferred chromosomal clonal trees. (f) Distribution of observed VAFs for the top five most frequently occurring latent VAFs (indicated with a dashed line). (g) Inferred clonal tree for chromosome 3, with labels for introduced CNAs and number of introduced SNVs, as well as the cell clustering showing the number of assigned cells for each node. (h) Number of SNVs for each of the *k* = 4 SNV clusters, colored by the chromosome in which they occur. (i) CMB distribution for the set of SNVs introduced at clade A, which delineates clones A and I in the inferred Pharming clonal tree of chromosome 3. (j) UMAP projection of the cells colored by Pharming cell assignment in clonal tree of chromosome 3.

Tumor TN1 had nearly double the number of segments (*r* = 335) with more copy number states (mean: 9.2 for TN1 vs. 3.7 for TN3), posing a scalability challenge for Pharming and preventing inference of an integrated clonal tree for the entire genome (Fig. S12). As such, we ran Pharming on each of the 22 autosomal chromosomes (median of 13.5 segments and 434 SNVs per chromosome, see Fig. 4c). To maintain consistency, we utilized shared DCFs δ to obtain chromosome-specific clonal trees and cell clusterings (*T* ^(1)^, *ϕ*^(1)^), …, (*T* ^(22)^, *ϕ*^(22)^) with the same underlying SNV cluster tree 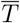. We repeated this process for *k* ∈ {4, 5, 6} SNV clusters. For each number *k* of SNV clusters, we selected the SNV cluster tree 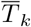 with *k* clusters such that the total cost 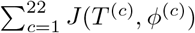 across all clonal trees was minimum. Finally, following BIC analysis, we arrived at the final set of *k* = 4 SNV clusters and corresponding SNV cluster tree 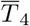 and inferred chromosomal clonal trees (Fig. 4d, Fig. S14). Fig. 4e shows the distribution of the CMB for each chromosome aggregated over the clades where the *k* = 4 SNV clusters were introduced. We observed a high median CMB of 0.36 for cells within the clade and a median of 0 for cells assigned outside clades of introduced SNVs. This suggests that Pharming is able to effectively use the SNV signal to partition cells in the clonal trees.

Next, we assessed the genotypes *g* and corresponding latent VAFs *f* inferred by Pharming (Def. 2). The sparsity of the coverage prevents us from directly comparing the observed VAF *a*_*i,j*_*/d*_*i,j*_ and latent VAF (*f* (*ϕ*(*i*), *j*) for a cell *i* and SNV *j*. Therefore, inspired by the approach taken in CNAqc [27], we pooled nodes in the clonal tree that have the same latent VAF for each SNV *j*, followed by aggregating the read counts of all cells *N* assigned to these nodes. This facilitated a calculation for the observed VAF, i.e., 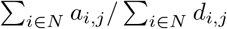 that is a better estimator of the latent VAF given a clonal tree. We restricted our analysis to only those SNVs with at least 15 total pooled reads, showing the observed VAFs for the top five most frequently occurring latent VAFs in Fig 4f. We expected that the peak of the observed VAF distribution would occur at the underlying latent VAF, which was indeed the case for the most frequent latent VAFs for this tumor as well as for TN3 (Fig. S13d). However, we also observed a secondary peak for latent VAF 1*/*4 that is centered at 1*/*2 as well as a small secondary peak at 1*/*4 for the latent VAF of 1*/*2. As these tumors were previously reported to have undergone whole genome doubling [14], these results suggest that Pharming may struggle to disambiguate whether the SNV occurred before or after genome doubling when these events occur within the trunk of the clonal tree. We also noted that that the SNVs comprising each SNV cluster *q* come from segments and chromosomes across the entire genome (Fig 4h) as opposed to clustering by chromosome. Furthermore, these SNV cluster assignments are well supported by both CMB (Fig. 4e) and the pooled observed VAF (Fig 4f).

Lastly, we focused our analysis on a single clonal tree for chromosome 3 comprised of 19 segments and 703 SNVs (Fig. 4g, Fig. S14). The majority of these SNVs (492) occurred on the trunk of the tree. Additionally, Pharming indicated genome doubling events in 11 segments, spanning 101.4 Mb (53.6%) of the chromosome. Moreover, these doubling events occurred in adjacent segments despite the fact that Pharming does not consider dependencies between nearby segments. The remaining SNVs comprised two subclonal SNV clusters consisting of 27 and 184 SNVs. The 184 SNVs comprising the largest subclonal cluster are well supported by the CMB metric (Fig. 4i, Fig. S14). Interestingly, based on the Pharming clonal tree, the cells in node I and the the cells in node A are only differentiated by the introduction of these 184 SNVs and no copy number changes. Indeed, we validated that these cells would appear clustered together in a low dimensional embedding of the allele-specific copy number profiles (Fig 4j, Fig. S14). The fact that Pharming is able to differentiate these clones using SNVs is noteworthy because the commonly used baseline approach clustered these cells together (Fig. S14a), potentially obscuring key events in the evolutionary history of the tumor. See Fig. S14 and Fig. S13 for additional results.

### 3.3 Ovarian cancer cell lines

We assessed the performance of Pharming on a dataset comprised of three clonally related ovarian cancer cell lines sequenced with DLP+ [13]. This dataset contained *n* = 890 cells and *m* = 13,428 SNVs within *r* = 409 segments obtained from CNRein [20]. Due to the high number of segments, we opted to run Pharming individually on each of the three cell lines (SA922 with 220 cells, SA1090 with 426 cells, SA921 with 244 cells) on a random downsampling of segments across the genome — see Appendix B.4. As detailed above, we used the CMB metric to validate the inferred clonal tree for each cell line, finding similarly strong performance on CMB for both clonal and subclonal SNV clusters, comparable to our observations with the breast cancer tumors (Fig. S15, Fig. S17, Fig S16). Critically, cell lines SA922 and SA921 were both previously hypothesized to have undergone a whole genome duplication [13, 28], and this is indeed reflected in the Pharming inferred clonal trees (Figs. S15, S17) while absent from the inferred clonal tree for SA1090 (Fig. S17). Although analyzing this data on all 890 cells and 409 segments simultaneously would be ideal, this analysis does demonstrate that our method is effective on datasets with a small number of cells and SNVs, despite the low sequencing coverage.

## 4 Discussion

In this work, we introduced the Joint Clonal Tree Inference (JCTI) problem for inferring a joint CNA and SNV clonal tree from low pass scDNA-seq, and proposed a heuristic algorithm, Pharming, to solve it. We demonstrated the accuracy of our algorithm on both *in silico* experiments and real data consisting of two breast cancer tumors sequenced with ACT [14] and three clonally related cell lines sequenced with DLP+ [13]. On *in silico* experiments, we showed that Pharming faithfully reconstructs the evolutionary history of CNAs and SNVs with respect to a known ground truth, as well as placement of cells onto the clonal tree. On experimental data, we showed that Pharming provides accurate and high-resolution joint reconstruction of CNA and SNV evolution.

There are several directions for future work. First, the scalability issues can be overcome in a number of ways, including re-implementing Pharming solely in C++ as well as parallelizing our implementation. Some of the scalability challenges could also be overcome by first using Phertilizer to identify the major clades of the clonal tree and then using Pharming to get a high-resolution joint CNA and SNV view of each clade. Second, Pharming could be extended to support parallel evolution of copy number states, i.e., violations of the infinite alleles assumption. Potential violations may be identified by looking at segments that do not integrate well with the other segments during progressive integration. Once identified, segments could be subjected to a more detailed inference by doubling one or more copy states before enumerating candidate CNA trees. Third, our progressive integration can be made more aware of dependencies between segments. For example, copy number events that span multiple segments should imply a consistent CNA evolutionary history within each segment. Fourth, the observed VAF distributions obtained by pooling cells with the same assigned latent VAF (Fig. 4f) highlight an opportunity to improve Pharming genotype inference of SNVs. In particular, one could alter the objective function *J*(*T, ϕ*) to asses the read count data of clones rather than individual cells with respect to each clone’s inferred latent VAF. Finally, our algorithmic approach is easily extendable to bulk DNA sequencing data or even joint bulk and single-cell data with only minor adjustments required.

---

**Table S1:**
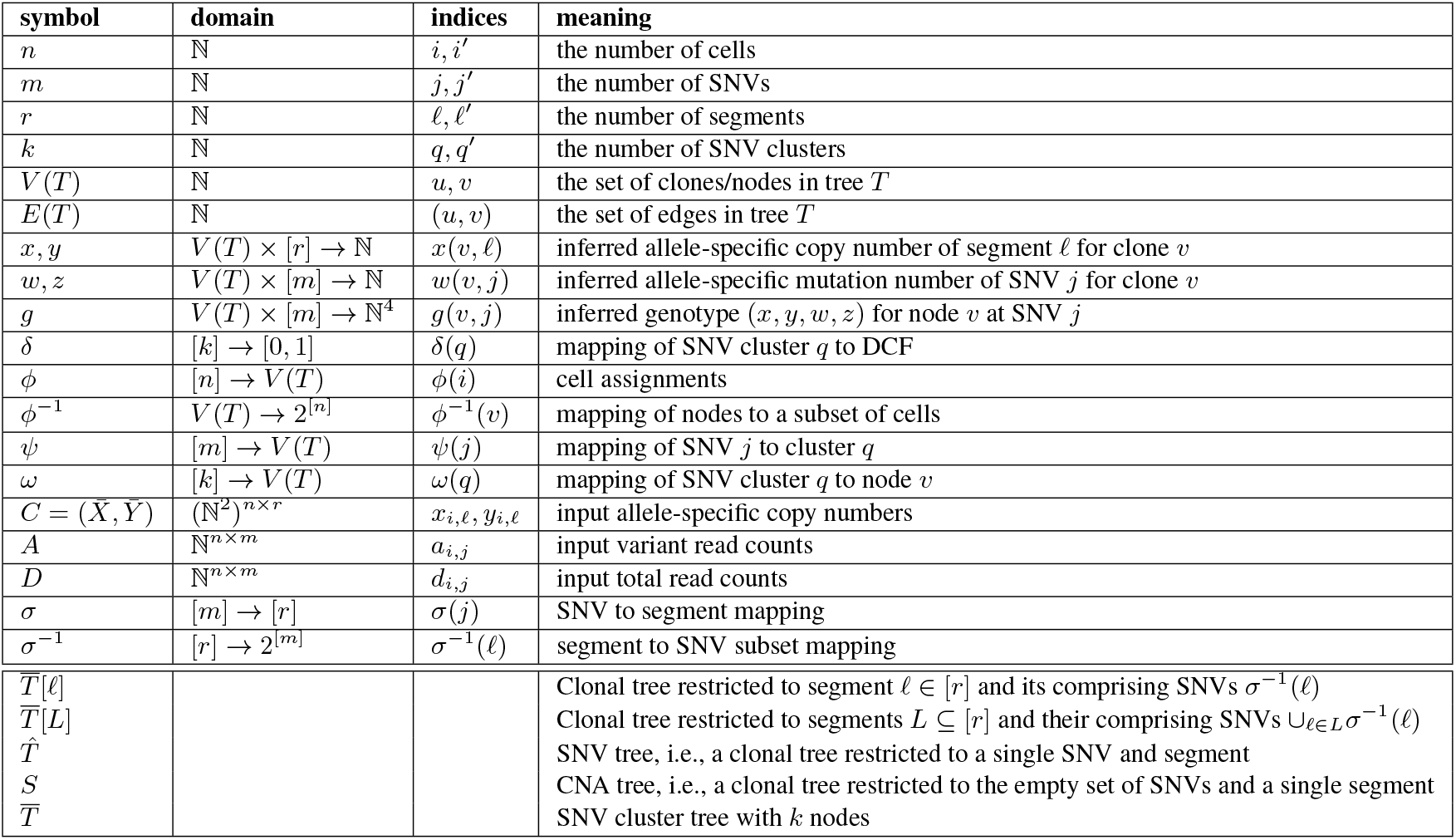
Summary of notation.

## A Supplementary Methods

### A.1 Pharming implementation of DCF clustering

A key input to Pharming is *k* DCFs. To obtain these values, one may run existing algorithms, such as DeCiFer [25]. However, for ease of use, Pharming includes its own re-implementation of this algorithm with optional modifications.

In particular, the Pharming implementation optionally enforces that every SNV tree assignment is consistent with a CNA tree *S*[*ℓ*] for all *m*_*ℓ*_ = *σ*^−1^(*ℓ*) SNVs occurring within segment *ℓ*. This modification works by adding a step to select a CNA tree *S* for each segment *ℓ*. During the SNV cluster and SNV tree assignment step of the DeCiFer algorithm, we first enumerate the set 𝒮 [*ℓ*] of all CNA trees for the given copy number states in each segment *ℓ*. Then for each CNA tree *S*, we enumerate the set of valid SNV trees 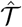, such that each SNV tree 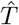 is consistent with CNA tree *S*. SNVs in segment *ℓ* may only be assigned to an SNV tree in the set 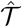. We use the DeCiFer posterior probability to select a CNA tree *S*[*ℓ*] ∈ 𝒮 [*ℓ*] and a corresponding assignment of SNVs to valid SNV trees for all segments prior to moving on to the cluster center optimization stage of the DeCiFer algorithm.

In practice, the number of inferred segments by copy number callers may be large, i.e., *r >* 100, and our proposed modification increases the running time by a factor of *r*. In these cases, our implementation offers a speed up by randomly sampling a user-specified number of segments during each restart. This is a valid approach because the underlying assumption is that the SNVs will cluster into a small number *k* of clusters across the genome. Therefore, only a small number of segments may be needed to detect the presence of an SNV cluster. It also allows us to quickly prune poor random initialization values of the DCFs δ. After running our DCF clustering algorithm for a specified number of restarts, we return the top *n* inferred DCFs δ with the highest posterior probability and use these to initialize restarts of our DCF clustering for all *r* segments.

### A.2 Inferring a clonal tree for a single segment

A tree *T* [*ℓ*] is a clonal tree for segment *ℓ* such that the genotypes *g*(*v, j*) are restricted to the set *σ*^−1^(*ℓ*) ⊆ [*m*] of SNVs located in segment *ℓ*. Since we focus on a single segment *ℓ*, we expect the size of the set 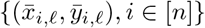 depicting the unique allele-specific copy number states with proportion of cells having that state occurring in segment *ℓ* to be small. Consequently, it is feasible to enumerate a set 𝒮 [*ℓ*] of CNA trees (Def. 3) with each node labeled by exactly one CNA state in this set the root node labeled with copy number (1, 1). Optionally, a user-specified threshold on the minimum proportion of cells that support a given copy number state within segment *ℓ* may be provided to reduce the size of set 𝒮 [*ℓ*] of CNA trees that are considered. As discussed in the main text, the Segment Tree Inference(STI) problem is a variant of the Joint Clonal Tree Inference, which focuses only on a single segment *ℓ* for a fixed SNV cluster tree 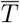 and CNA tree 𝒮 ∈ [*ℓ*]. We now describe how Pharming solves the STI problem.

**Figure S1:**
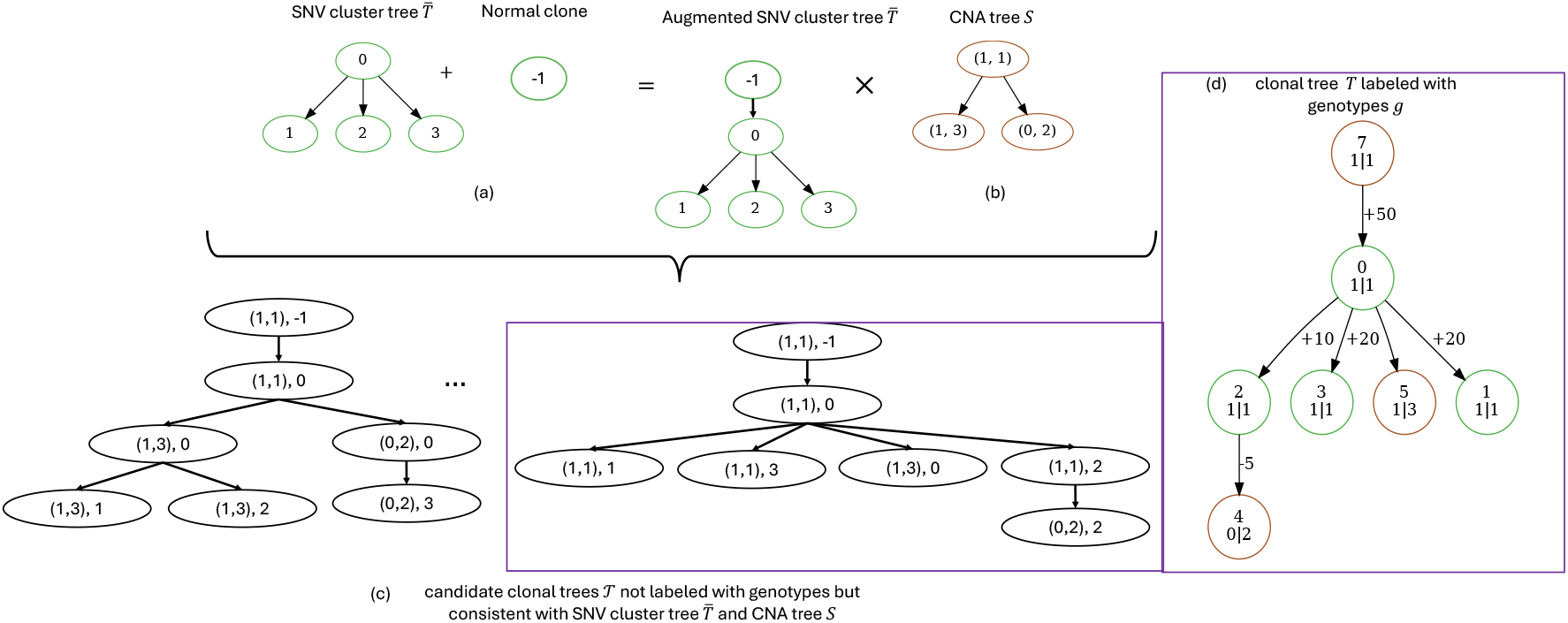
Enumerating clonal trees from a given SNV cluster tree 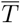 and CNA tree *S*. (a) An SNV cluster tree 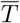 augmented with a normal clone as the root. (b) A example CNA tree *S*. (c) An example of enumerating candidate clonal tree such that each clonal tree *T* ∈ 𝒯 is consistent with both the SNV cluster tree 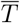 —see (a) and CNA tree *S* — see (b). These candidate clonal trees are not yet labeled by genotypes. (d) A clonal tree *T* that is labeled by genotypes, i.e., SNVs are assigned to an SNV cluster and an SNV tree and corresponds to the last clonal tree in the set — see (c).

We start by augmenting the SNV cluster 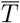 by attaching a root node to 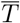, representing a normal clone harboring no SNVs, i.e, 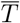 (Fig. S1a). Next we enumerate all clonal trees 𝒯 [*ℓ*] consistent with the CNA tree *S* and the SNV cluster tree 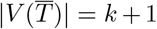 by constructing a product graph of *S* and 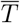 and applying the Gabow-Myers algorithm [29]. Each clonal tree *T* [*ℓ*] that is consistent with with both *S* and 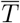 is retained (Fig. S1c). To then solve the SIT problem for each retained tree, we infer the optimized genotypes *g* and cell assignment *ϕ* for each retained clonal tree *T* [*ℓ*] (Fig. S1)d). This is achieved via a coordinate descent algorithm consisting of three phases: (i) initialize the genotypes, (i) optimize cell assignment *ϕ* given the clonal tree *T* [*ℓ*] with fixed genotypes *g* and (iii) optimize the genotypes *g* given a fixed cell assignment *ϕ*. Steps (ii) and (iii) are repeated until the objective *J*(*T* [*ℓ*], *ϕ*) converges or a maximum number of iterations is reached. Next, we provide additional details on each of these phases.

#### Initialization of genotypes

Given a clonal tree *T* [*ℓ*] not labeled by genotypes but indicating the evolutionary relationships of SNV clusters to and copy number states for segment *ℓ*, we identify for each SNV cluster *q*, the set of valid SNV trees 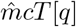 such that the clonal tree is consistent with each SNV tree in this set for an SNV placed in cluster *q*. Now we initialize genotypes *g* by assigning each SNV *j* to a cluster *q*, i.e., *ψ*(*j*) = *q* and an SNV tree 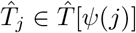.

Note that 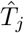 is a clonal tree for a single SNV. Therefore, we compute the cost 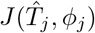 by greedily assigning cells to nodes in clonal tree 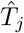 — see below for a details on optimizing cell assignments. We compute this cost for each SNV cluster *q* and SNV *j* and retain the set *Q* of SNV clusters which contains an SNV tree 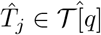 with minimum cost.

In words, based on the the clonal *T* [*ℓ*], two SNV clusters *q* and *q*^*′*^, may have identical sets of SNV trees, i.e., 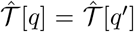 if both SNV clusters *q* and *q*^*′*^ have the same evolutionary relationships to the copy number states. For example, cluster *q* and *q*^*′*^ may both be introduced after the copy number state 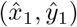 but before states 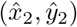 is obtained with cluster *q* being ancestral to cluster *q*^*′*^ in clonal tree *T* [*ℓ*]. In this case, we break ties between cluster assignments by utilized the posterior probability of the DCFs corresponding to each cluster *q* in the set *Q*. See Satas et al. [25] for details and derivations of the posterior probability. Following the Satas *et al*. [25] probabilistic model for DCFs, we use a binomial model to compute the following likelihood

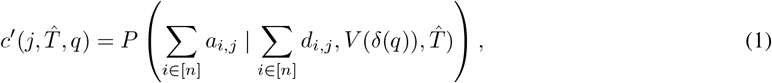

where 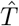 is the SNV tree 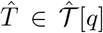 that has minimum cost for SNV cluster *q* ∈ *Q* and *V* (*d*) is the function that converts DCF *d* to a VAF utilizing SNV tree 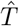 [25]. We then select the SNV cluster *q*∈ *Q* and corresponding SNV tree 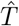 with minimum cost 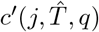.Finally, given an SNV *j*, SNV tree 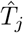 and SNV cluster assignment *ψ*(*j*), the genotype *g*(*v, j*) of each node *v* ∈ *V* (*T* [*ℓ*]) is uniquely determined.

#### Optimizing cell assignments

Given a clonal tree *T* [*ℓ*] for segment *ℓ* and genotypes *g*, we then seek the cell assignment *ϕ* that minimizes *J*(*T* [*ℓ*], *ϕ*). This is accomplished by independently finding a optimal assignment of each cell *i* to a node *u* in clonal tree *T* [*ℓ*], i.e., cell, *ϕ*(*i*) = arg min_*u*∈ *V* (*T* [*ℓ*])_ *J*(*T* [*ℓ*], *u*), where we abuse notation slightly such that *J*(*T* [*ℓ*], *u*) is the cost of assigning cell *i* to node *u* in clonal tree *T* [*ℓ*].

#### Optimizing genotypes

Given a clonal tree *T* [*ℓ*] for segment *ℓ* and a fixed cell assignment *ϕ*, we then infer genotypes *g* that optimizes *J*(*T* [*ℓ*], *ϕ*). This is accomplished by independently computing the cost of assigning an SNV *j* to each SNV cluster *q* and SNV tree 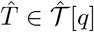. For each SNV *j*, we select the SNV cluster *q*^∗^ and SNV tree 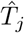 that yield the genotypes *g*(:, *j*) across all nodes in clonal tree *T* [*ℓ*] that minimizes *J*(*T* [*ℓ*], *ϕ*) for SNV *j*. In the event of ties between SNV clusters, we prefer the SNV cluster closest to the root of the tree. In the event of ties for SNV tree assignments within a cluster, we prefer SNV trees without indicated loss of SNV.

##### A.2.1 Model selection

The key hyperparameter for Pharming is the number *k* of mutation clusters in the underlying SNV cluster tree *T*. This parameter choice also influences the estimates of the input DCFs δ. To avoid overfitting the data, we propose a two-stage model selection process. In stage one, the DCF clustering algorithm of choice is run for a wide range of reasonable values of *k*. Initial model selection utilizing standard approaches for clustering, such as the elbow method or silhouette score, are then applied to narrow down the range of *k* values and corresponding DCFs δ given to Pharming for clonal tree inference. Set *K* denotes these prioritized choices for the number of mutation clusters used as a hyperparameter for Pharming.

In the second stage, we make use of the Bayesian information criterion (BIC), i.e., BIC(*k*), defined as a function of the number of SNV clusters *k*, the number of SNVs *m* and the likelihood of the variant reads *A* given total read counts *D*, a clonal tree *T* with *k* SNV clusters, cell assignment *ϕ* and a sequencing error probability.

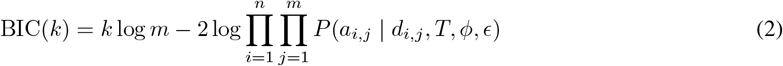

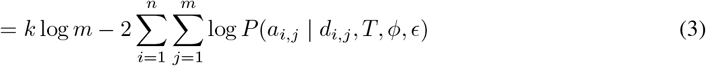

Pharming outputs the clonal tree *T* and cell assignment *ϕ* for the number *k*^∗^ = arg min_*k∈ K*_ BIC(*k*) of SNV clusters.

## B Supplementary Results

### B.1 Simulation details

#### B.1.1 Details on simulation method

We generated *in silico* scDNA-seq data modeling a heterogeneous tumor with both CNAs and SNVs. For each of the *r* segments, a CNA tree is sampled uniformly at random from a set of enumerated CNA trees given an input set of allowable combinations of CNA states. We then merge each CNA tree into a clonal tree via a random walk on the product graph of the clonal tree and CNA tree to be added. SNVs are randomly assigned to clusters and as well as segments by sampling from a Dirichlet distribution with symmetric concentration parameter 2. Next, SNV clusters are assigned to nodes uniformly at random with one SNV cluster assigned to a truncal node and at most one SNV cluster assigned to non-truncal nodes. Each SNV is phased at random and evolved down the clonal tree given its SNV cluster and segment assignment. Finally, the clone proportions at each node are sampled from a Dirichlet distribution, with rejection sampling enforce a minimum threshold on the smallest non-zero proportion.

To generate scDNA-seq variant and total read counts along with copy numbers for *n* cells, *m* SNVs and *r* segments, we randomly assign each of the *n* cells to nodes based on the sampled clonal proportions. We then draw total read counts from a Poisson distribution parameterized by the sequencing coverage and copy number and variant read counts are drawn from a beta-binomial distribution which introduces sequencing errors into our model for each of the *n* cells and *m* SNVS. Lastly, we introduce errors into the allele-specific copy numbers of the *in silico* cells with probability *ϵ* for each of the *n* cells and *r* segments. If errors are introduced for a cell and segment, we randomly select an allele and adjust the number profile of that allele by either one or two copies for to generate our observed copy number profiles.

#### B.1.2 Details on comparison methods

We compared against Phertilizer [5], which infers an SNV clonal tree aided by a low-dimensional cell embedding generated from copy number data. Here, we utilized the simulated allele-specific copy number profiles to generate a UMAP cell embedding [30]. Since Phertilizer outputs a cell assignment but not clonal CNA states of inferred nodes, we generated clonal CNA states by taking the consensus state of all cells mapped to each node *u* and each of the *r* segments.

We also benchmarked against a commonly used *ad hoc* method, which we previously referred to as the Baseline method [5]. Briefly, the Baseline method works by projecting the copy number profiles into a low dimensional embedding, i.e., UMAP, and then using a density-based clustering approach to obtain a final cell clustering. We then inferred clonal CNA states as described above for Phertilizer of each cell cluster and called the presence or absence for each SNV *j* by pooling read counts across the cell cluster to compute the observed variant allele frequency (VAF). We called an SNV present in a cell cluster if the observed VAF for SNV *j* was greater than 0.05. Next, we provided SCITE [7] with the observed presence or absence of each SNV *j* for each cell cluster to generate an output mutation tree with cell clusters mapped to nodes. Lastly, we collapsed SNVs that formed a linear chain in between branched nodes and/or nodes with mapped cells into SNV clusters. We refer to this method as Baseline+SCITE.

### B.2 Performance metrics

In this section, we provide details on the calculation of the performance metrics used for both simulated and real data analysis.

#### B.2.1 Simulation performance metrics

##### Cell placement accuracy

Cell placement accuracy is a measure to simultaneously assess the assignment of cells to clones as well as the accuracy of the inferred evolutionary relationships of those clones. More specifically, we compute the weighted average of the *ancestral* pair recall (APR), *incomparable* pair recall (IPR) and *clustered pair recall* (CPR) [2] (Fig. S2).

For any two cells *i* ≠ *i*^*′*^ there are three possible placements in the tree: (i) *ancestral*, if *i* is assigned to a node that is distinct and ancestral to the node where *i*^*′*^ is assigned; (ii) *clustered*, if *i* and *i*^*′*^ are both assigned to the same node; and (iii) *incomparable*, if *i* and *i*^*′*^ are assigned to distinct nodes that occur on distinct branches of the tree. The APR assesses the ratio of ancestral pairs from *T*^∗^ recalled in *T*, whereas CPR and IPR do so for clustered and incomparable pairs, respectively. The weighted average of these recall scores is computed by weighting each pair recall score by the number of those pairs in the ground truth tree. Fig. S2 gives an example for this calculation.

##### CNA tree recall

For each segment *ℓ*, we construct the underlying CNA tree *S*[*ℓ*] from clonal tree *T* by removing all edges (*u, v*) from *T* whenever *x*(*u, ℓ*) = *x*(*v, ℓ*) and *y*(*u, ℓ*) = *y*(*v, ℓ*). The nodes of CNA tree *S*[*ℓ*] are then labeled by (*x*(*u*), *y*(*u*)) for each node *u*. Then,

**Figure S2:**
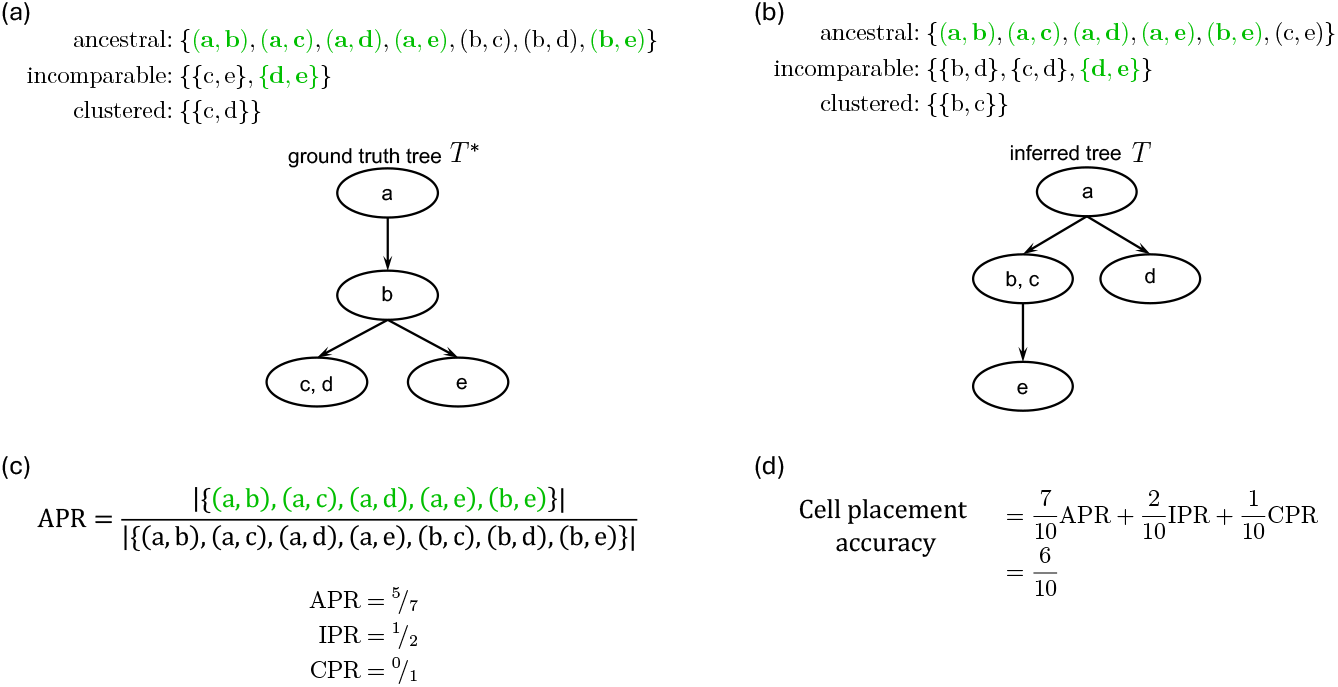
Example computation of the *Cell placement accuracy* performance metric. (a) The cell pair sets for *ancestral, incomparable, clustered* for the ground truth tree *T*^∗^. (b) The cell pair sets for the inferred tree *T*. (c) An example computation of the *ancestral* pair recall metric with corresponding values of IPR and CPR. (d) An example computation of the *cell placement accuracy*.

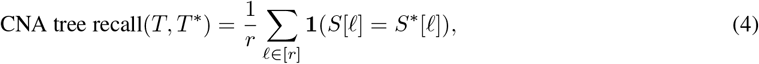

where *T* is the inferred clonal tree and *T*^∗^ is the ground truth.

**Figure S3:**
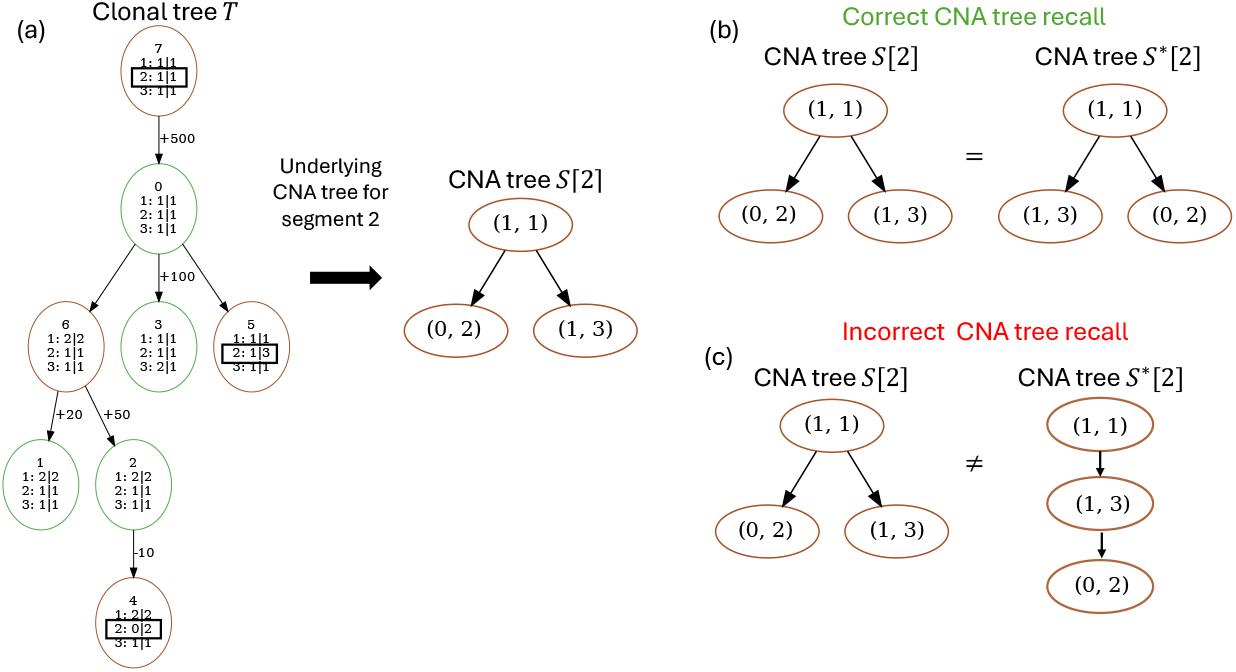
Example computation of the *CNA tree recall* performance metric. (a) Example of obtaining the underlying CNA tree *S*[2] for segment 2 from clonal tree *T*. Nodes with a change in copy number state for segment 2 are show are indicated with a boxes around the copy number state. (b) An example of correct CNA tree recall for segment 2. (c) An example of incorrect CNA tree recall for segment 2.

##### SNV tree recall

For an SNV *j* that occurs in segment *ℓ* with genotype 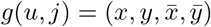 at node *u* in clonal tree *T*, we say the presence status 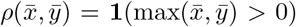.In words, we seek to modify the genotype of SNV *j* at node *u* to ignore both the allele-specificity and the number of mutated copies when considering the genotype of SNV *j* at node *u*. This allows us to compare the inferred evolutionary relationships between the CNA states and an SNV against methods that only consider presence or absence of SNV.

We construct the underlying SNV tree 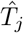 from clonal tree *T* by removing all edges (*u, v*) from *T* whenever *x*(*u, ℓ*) = *x*(*v, ℓ*), *y*(*u, ℓ*) = *y*(*v, ℓ*) and *ρ*(*u, j*) = *ρ*(*v, j*). In words, we remove edges from clonal tree *T* whenever the genotype of SNV *j* does not change when ignoring both allele-specificity and the number of mutated copies. The nodes of SNV tree 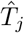 tree are then labeled by 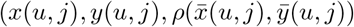 for each node *u*.

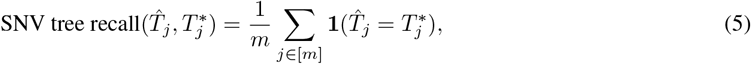

where 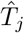 is the inferred SNV tree with modified genotype labels and 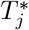 is the ground truth SNV tree with modified genotype labels.

**Figure S4:**
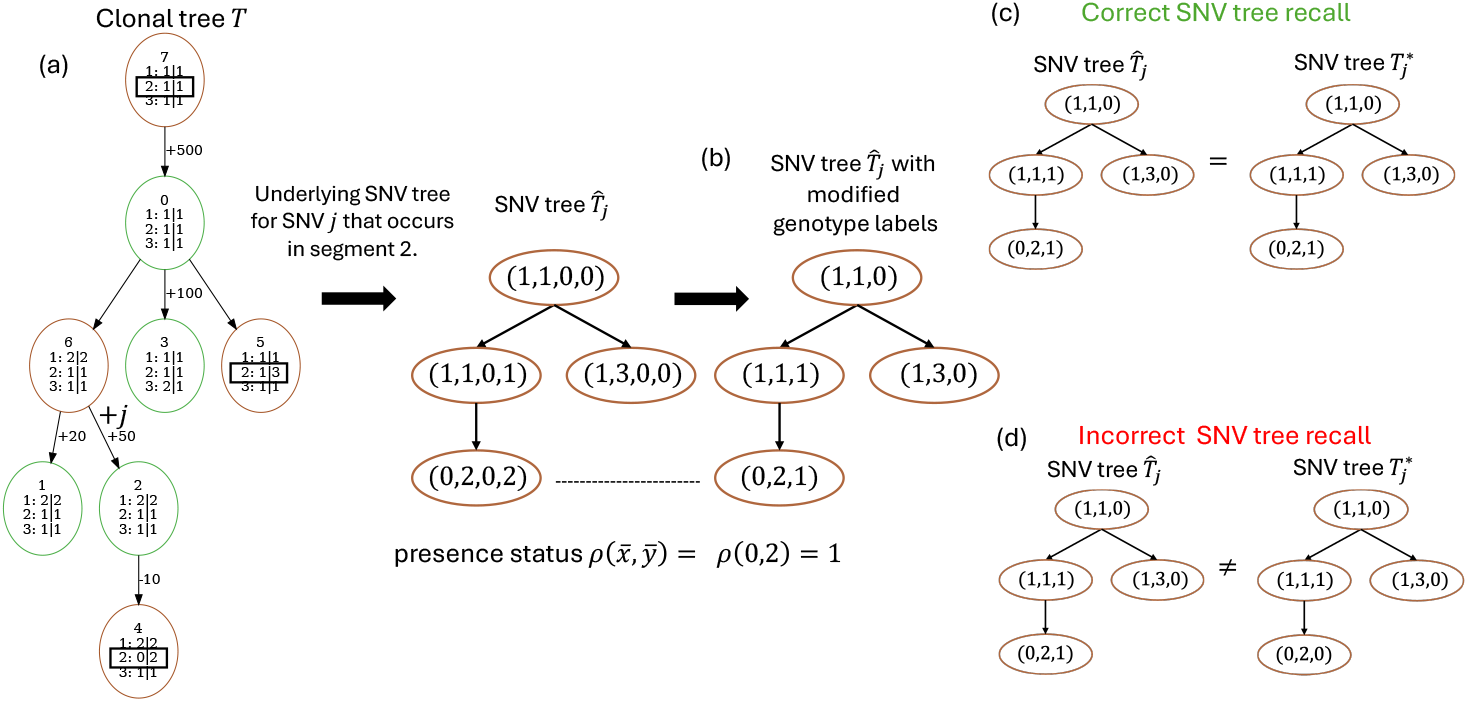
Example computation of the *SNV tree recall* performance metric. (a) Constructing the underlying SNV tree 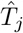 for SNV *j* in segment 2 from Clonal tree *T*. (b) Modification of the genotype labels for SNV tree 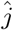 to only consider presence or absence of SNV *j*. (c) An example of a correctly inferred SNV tree 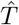. (d) An example of an incorrectly inferred SNV tree 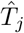.

##### Genotype similarity

Correctly genotyping a cell for each of the *m* SNVs requires that the correct evolutionary relationships between the clones are inferred, the accurate assignment of cells to clones, or nodes in the clonal tree, and that for each SNV *j*, the correctly underlying SNV tree is inferred. Thus, we develop the genotype similarity metric to summarize overall performance of a solution to the Joint Clonal Tree Inference problem.

Given a clonal tree *T*, a cell assignment *ϕ* constructed from *n* cells and *m* SNVs, we construct an *n* ×*m* genotype matrix *G* = [*g*_*ij*_] where *g*_*ij*_ = (*x*(*ϕ*^∗^(*i*), *y*(*ϕ*^∗^(*i*), *ϕ*^∗^(*i*) + *z*(*ϕ*^∗^(*i*)). In words, for each cell *i*, we identify the node in the clonal tree to which is is assigned, i.e., *ϕ*(*i*), and then identify the genotype of SNV *j* at node *ϕ*(*i*), ignoring allele-specificity but considering the number of mutated copies.

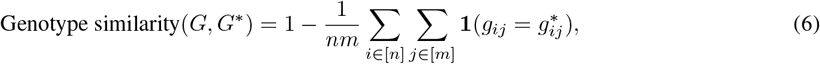

where *G* is the matrix of inferred genotypes and *G*^∗^ is the matrix of ground truth genotypes.

##### SNV loss recall, precision and F1

Since Pharming utilizes the Dollo evolutionary model, we assess how accurately it is able to identify loss of an SNV. Specifically, given a clonal tree *T*, let the set *M* be SNVs that are lost anywhere in the clonal tree. That is, there exists an edge (*u, v*) ∈ *E*(*T*^∗^) such that *w*(*u, j*) + *z*(*u, j*) *>* 0 and *w*(*v, j*) + *z*(*v, j*) = 0 for each *j*∈ *M*.

**Figure S5:**
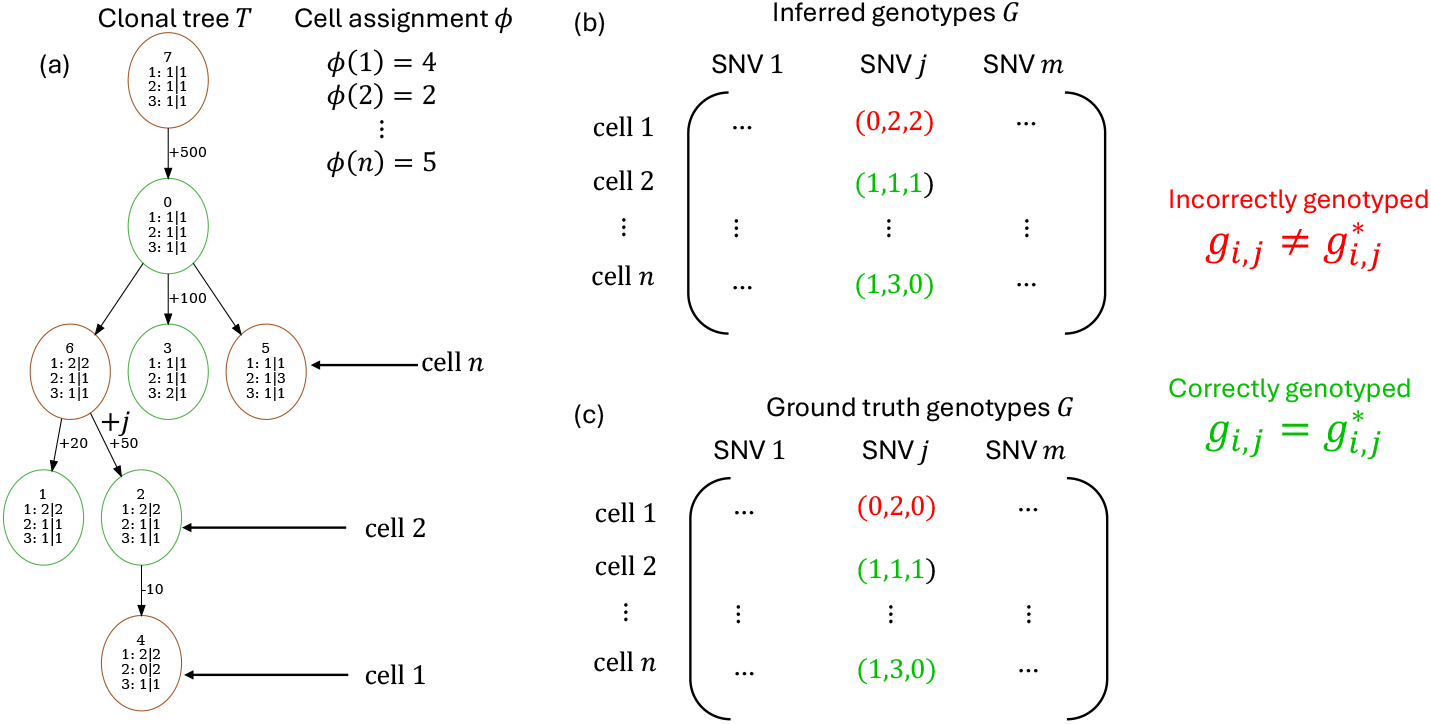
Example computation of the *Genotype similarity* performance metric. (a) An example clonal tree *T* and the corresponding cell assignment *ϕ*. (b) The genotype matrix *G* for the inferred clonal tree *T* and cell assignment *ϕ*. (c)The genotype matrix *G*^∗^ for the ground truth clonal tree *T*^∗^ and cell assignment *ϕ*^∗^. Genotypes that are correctly inferred are shown in green while incorrectly inferred ones are shown in red for SNV *j*.

We define the following loss metrics:

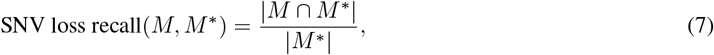

and

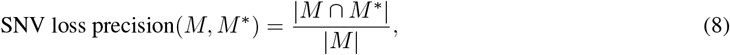

where set *M* are SNVs lost in the inferred clonal tree and set *M*^∗^ are SNVs lost in the ground truth clonal tree.

Lastly, we compute SNV loss *F* 1 as the harmonic mean of SNV loss recall and SNV loss precision, defined as follows.

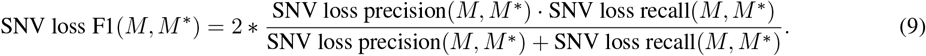

#### B.2.2 Real data performance metrics

##### Cell mutational burden (CMB)

To assess the quality of each inferred clade for real data, we previously developed a performance metric called *cell mutational burden* (CMB) [5], defined as

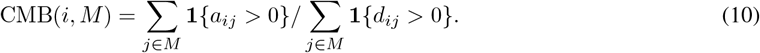

In words, CMB(*i, M*) is the fraction of mapped SNV loci *M* with mapped variant reads in cell *i*.

For a specified *clade u* or subtree rooted at node *u*, SNVs *M* are the SNVs gained at node *u* and never lost. More specifically, we exclude any SNV inferred as lost from this analysis.

CMB is designed to assess the goodness of fit of a proposed clonal tree without a known ground truth tree and succinctly captures the relationship between the inferred tree, clonal genotyping, and cell clustering. For a cell *i* placed within clade *u*, we expect CMB(*i, M*) to be high, although the value will depend on copy number. By contrast, for cells placed outside of clade *u*, we expect CMB(*i, M*) to be low.

### B.3 Data preprocessing

#### B.3.1 Processing the triple negative breast cancer tumors sequencing data from ACT

We merged reads from all cells into a single FASTQ file for each sample, aligned the merged reads to the Human reference genome hg19 (GRCh37) using bowtie2 (v.2.4.4), and sorted by samtools (v.1.15) forming pseudo-bulk samples. We then get a set of SNVs by running Mutect2 [31] in tumor-only mode on each pseudo-bulk sample, followed by FilterMutectCalls in GATK.

##### B.4 Processing the ovarian cancer cell lines sequenced with DLP+

We utilized SNVs and the variant and total read counts previously called by Laks et al. [13]. We utilized CNRein [20] to perform segmentation and haplotype-specific copy number calling.

For each cell line, we first filtered out SNVs that did not have at least two variant reads across all cells in the sample. Next, we ran DeCiFer [25] independently on each cell line for values of *k* ∈ {2, …, 4}. After this, we randomly sampled *r* = 25 segments from each cell line amongst segments that had a minimum of 20 SNVs per segment. Finally, we ran Pharming on each cell line, for the selected segments, for each value of *k*, using BIC to select the best value of *k* and corresponding Pharming clonal tree and cell assignment.

We used UMAP coordinates obtained from the binned read count embedding, with projected cells colored by the Laks et al. [13] clustering (Fig. S15c) and by the Pharming clone assignment (Fig. S15d). Laks et al. [13] previously inferred two distinct clusters for these cells using a density based clustering approach.

**Figure S6:**
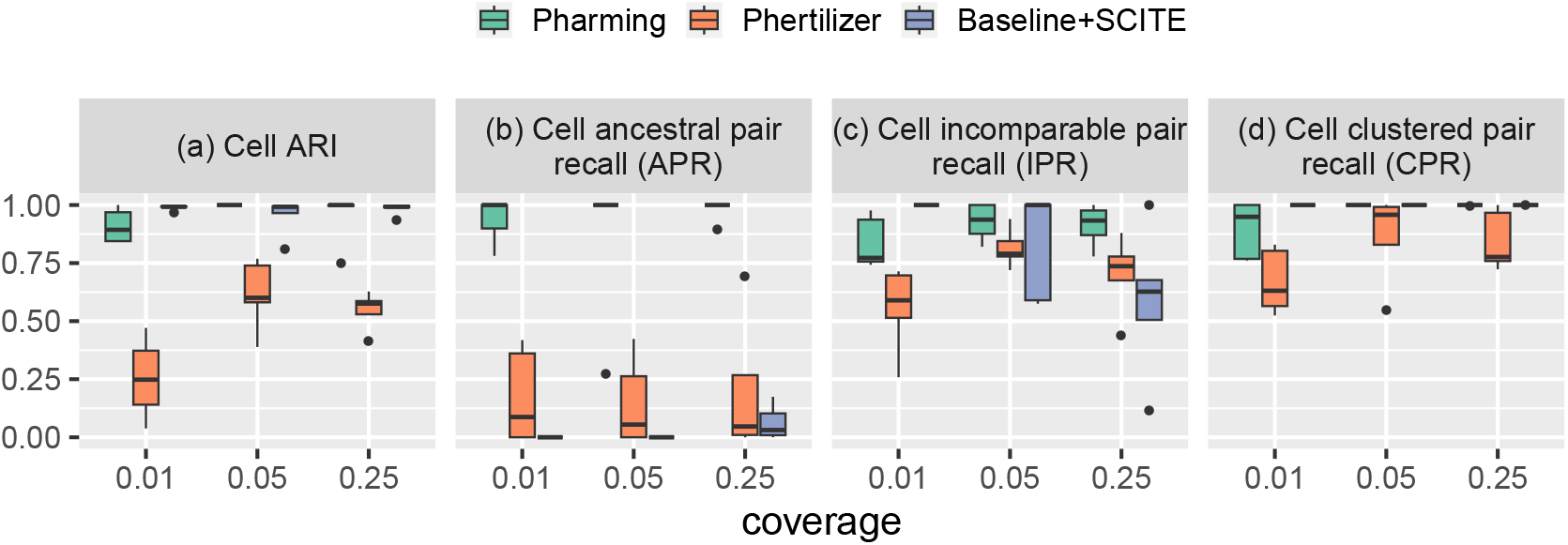
Benchmarking comparison of performance with known ground truth for cell clustering metrics with CNA error rate of 0. Results are shown for *in silico* experiments with 5000 SNVs, 1000 cells, CNA error rate 0,and coverage varying in 0.01×, 0.05×, 0.25× on the following metrics (a) *Cell ARI*, (b) *APR*, (c) *IPR*, and (d) *CPR*.

**Figure S7:**
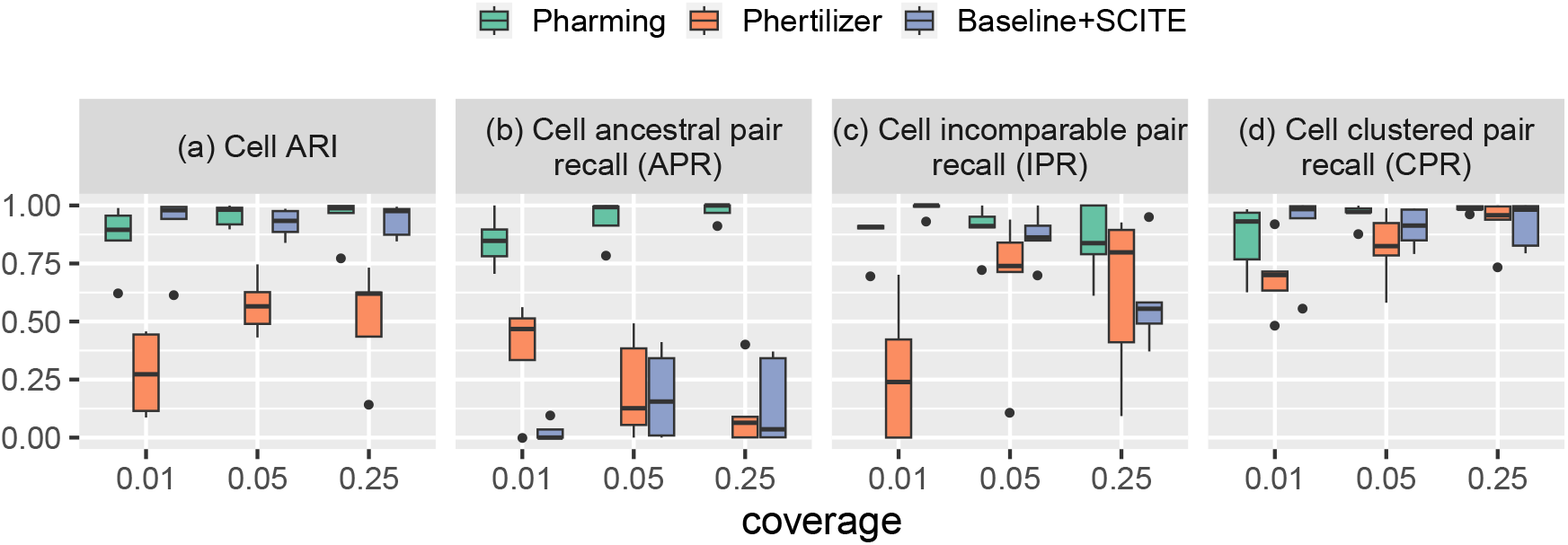
Benchmarking comparison of performance with known ground truth for cluster clustering metrics and CNA error rate of 0.035. Results are shown for *in silico* experiments with 5000 SNVs, 1000 cells, CNA error rate 0.035, and coverage varying in 0.01×, 0.05×, 0.25× on the following metrics (a) *Cell ARI*, (b) *APR*, (c) *IPR*, and *CPR*.

**Figure S8:**
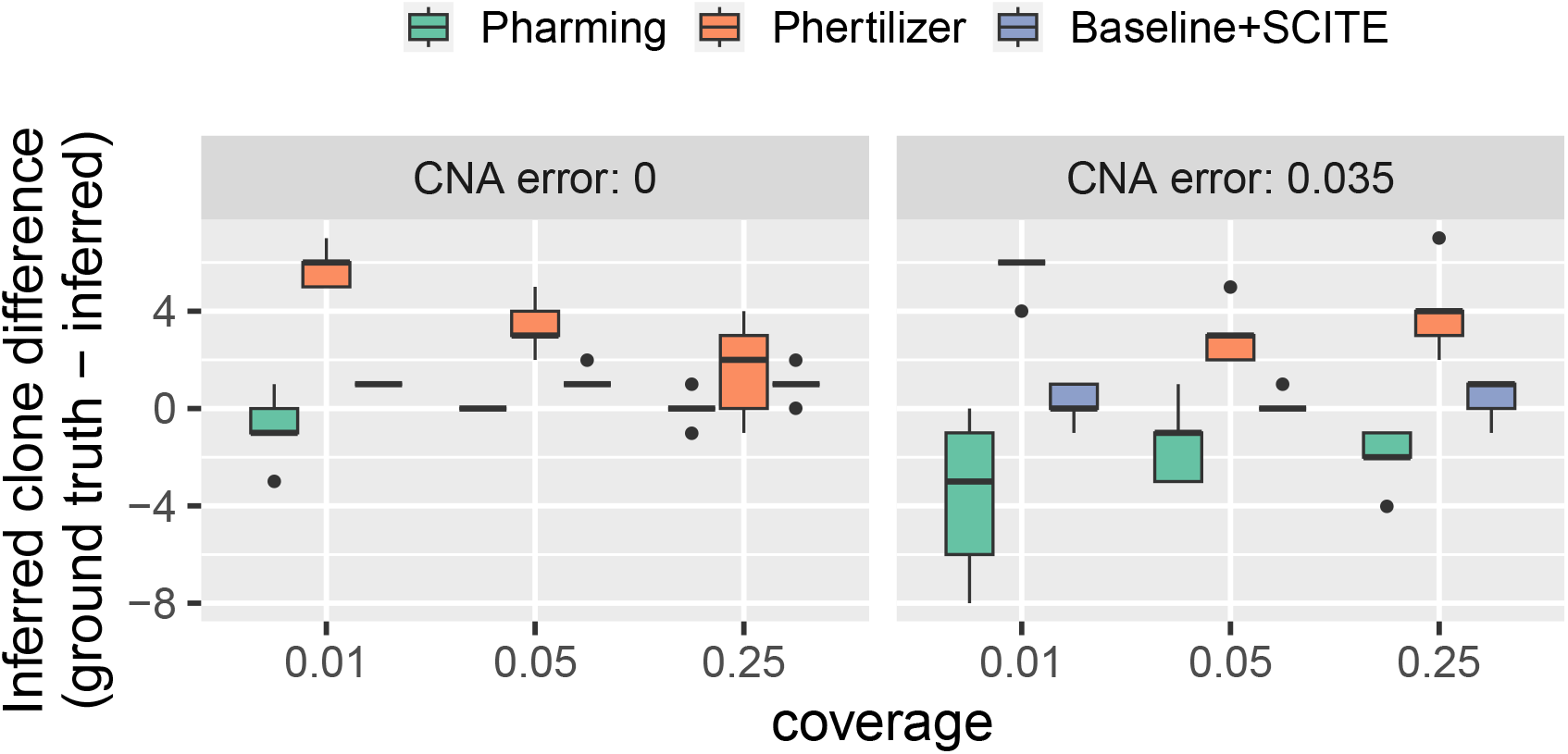
Benchmarking comparison of the difference between the number of ground truth clones and the number of inferred clones for *in silico* experiments. Results shown are with 5000 SNVs, 1000 cells, and coverage varying in 0.01×, 0.05×, 0.25×, for (a) CNA error rate 0 and (b) CNA error rate 0.035.

**Figure S9:**
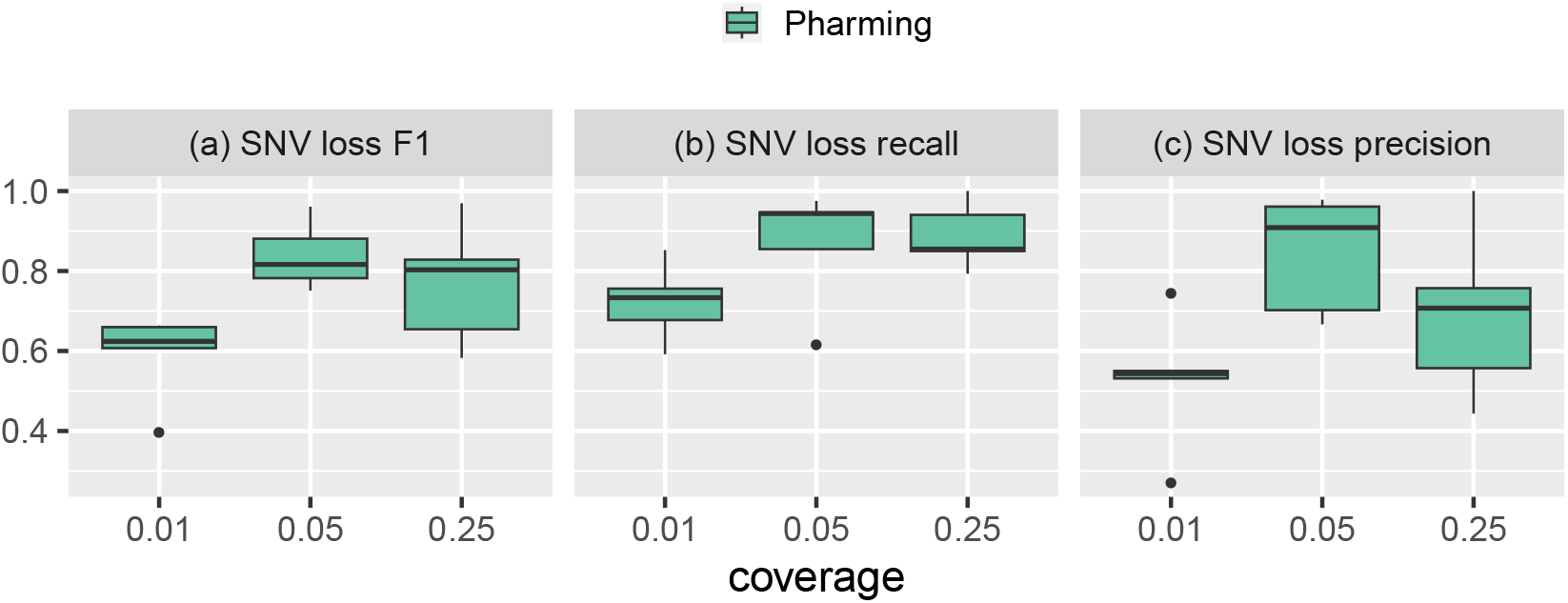
Pharming performance on SNV loss metrics with known ground truth and CNA error rate of 0. *Results are shown for in silico* experiments with 5000 SNVs, 1000 cells, CNA error rate 0, and coverage varying in 0.01×, 0.05×, 0.25× on the following metrics (a) SNV loss F1, (b) SNV loss recall, and (c) SNV loss precision.

**Figure S10:**
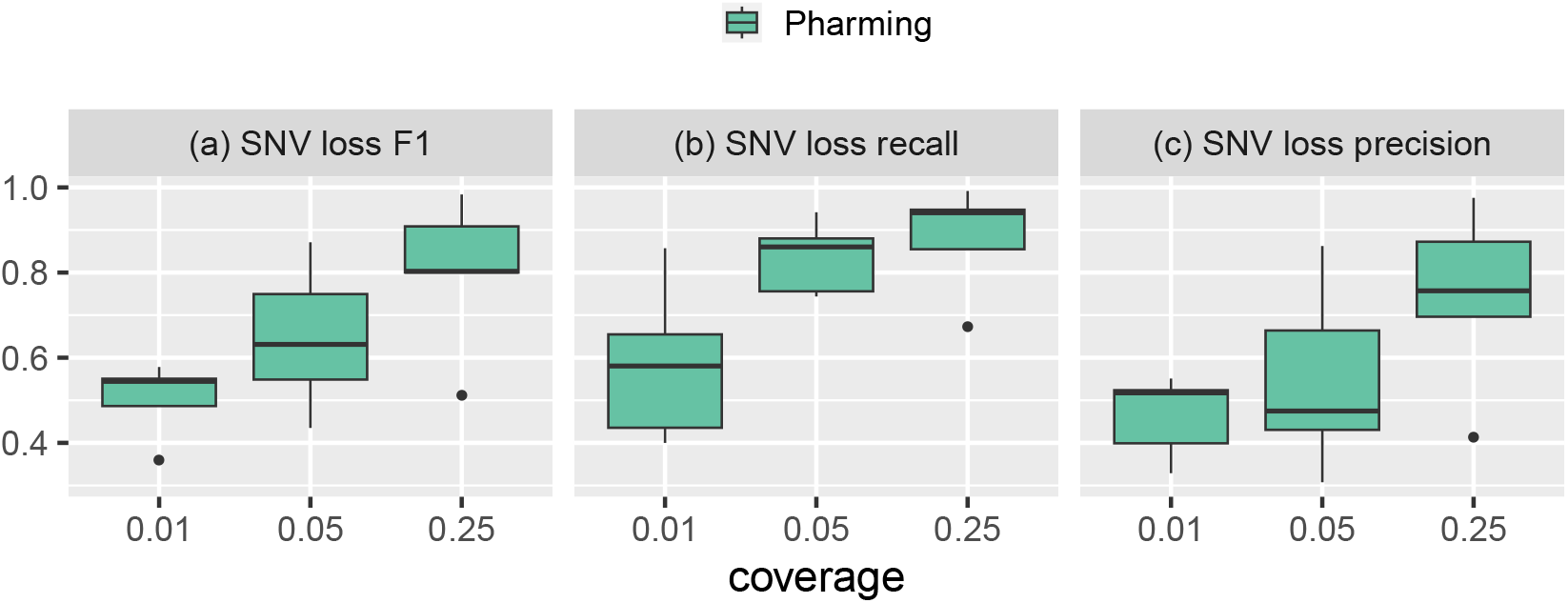
Pharming performance on SNV loss metrics with known ground truth with CNA error rate 0.035. Results are shown for *in silico* experiments with 5000 SNVs, 1000 cells, CNA error rate 0.035, and coverage varying in 0.01×, 0.05×, 0.25× on the following metrics (a) SNV loss F1, (b) SNV loss recall, and (c) SNV loss precision.

**Figure S11:**
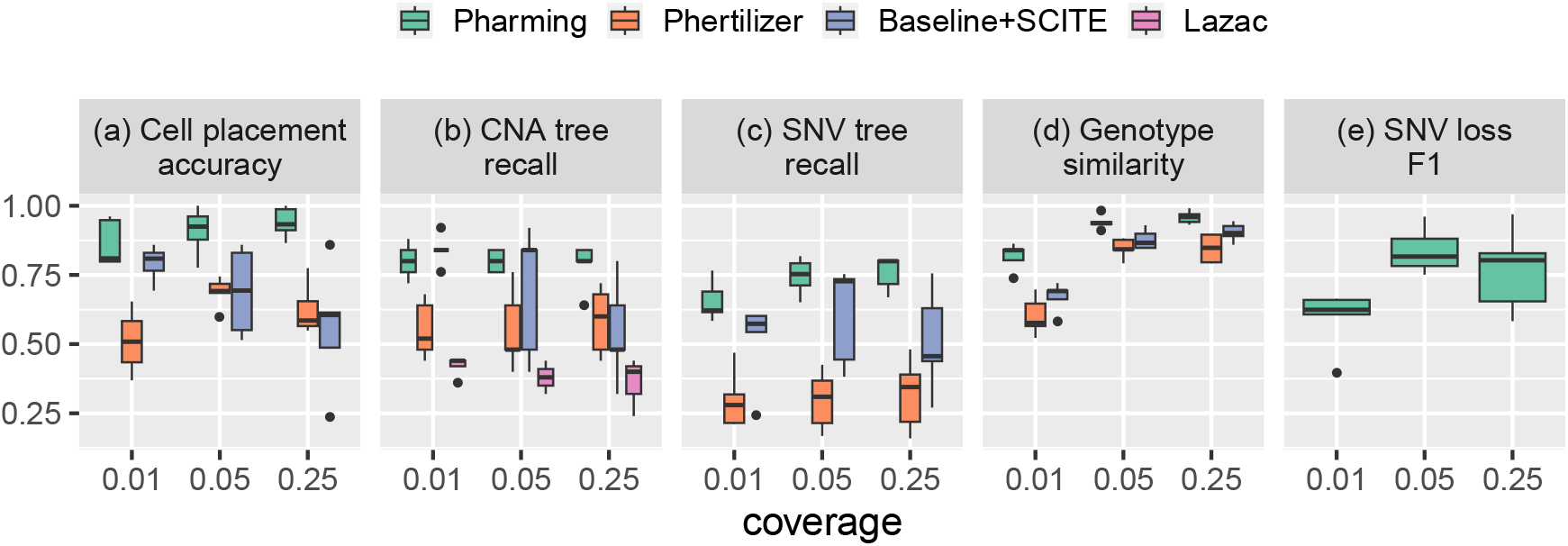
Benchmarking comparison of performance with known ground truth with CNA error rate of 0. Results are shown for *in silico* experiments with 5000 SNVs, 1000 cells, CNA error rate 0,and coverage varying in {0.01×, 0.05×, 0.25×} on the following metrics (a) Cell placement accuracy, (b) CNA tree recall, (c) SNV tree recall,(d) Genotype similarity and (e) SNV loss F1.

**Figure S12:**
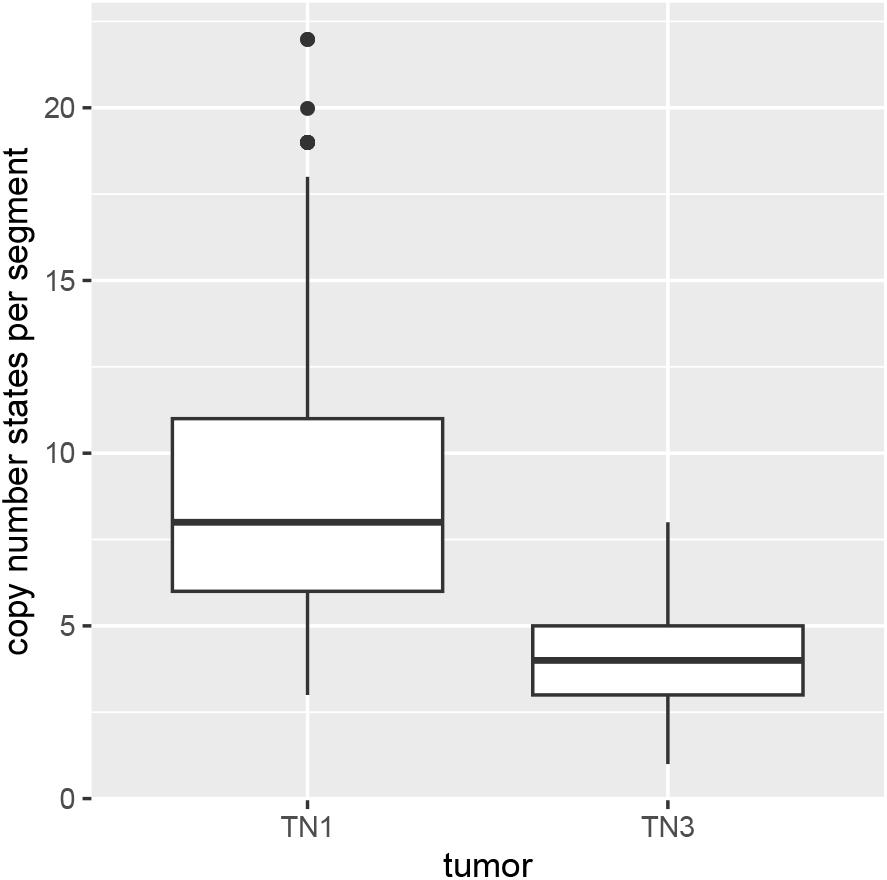
Distribution of the number of copy number states per segment from the input copy number profiles of the two triple negative breast cancer tumors TN1 and TN3.

**Figure S13:**
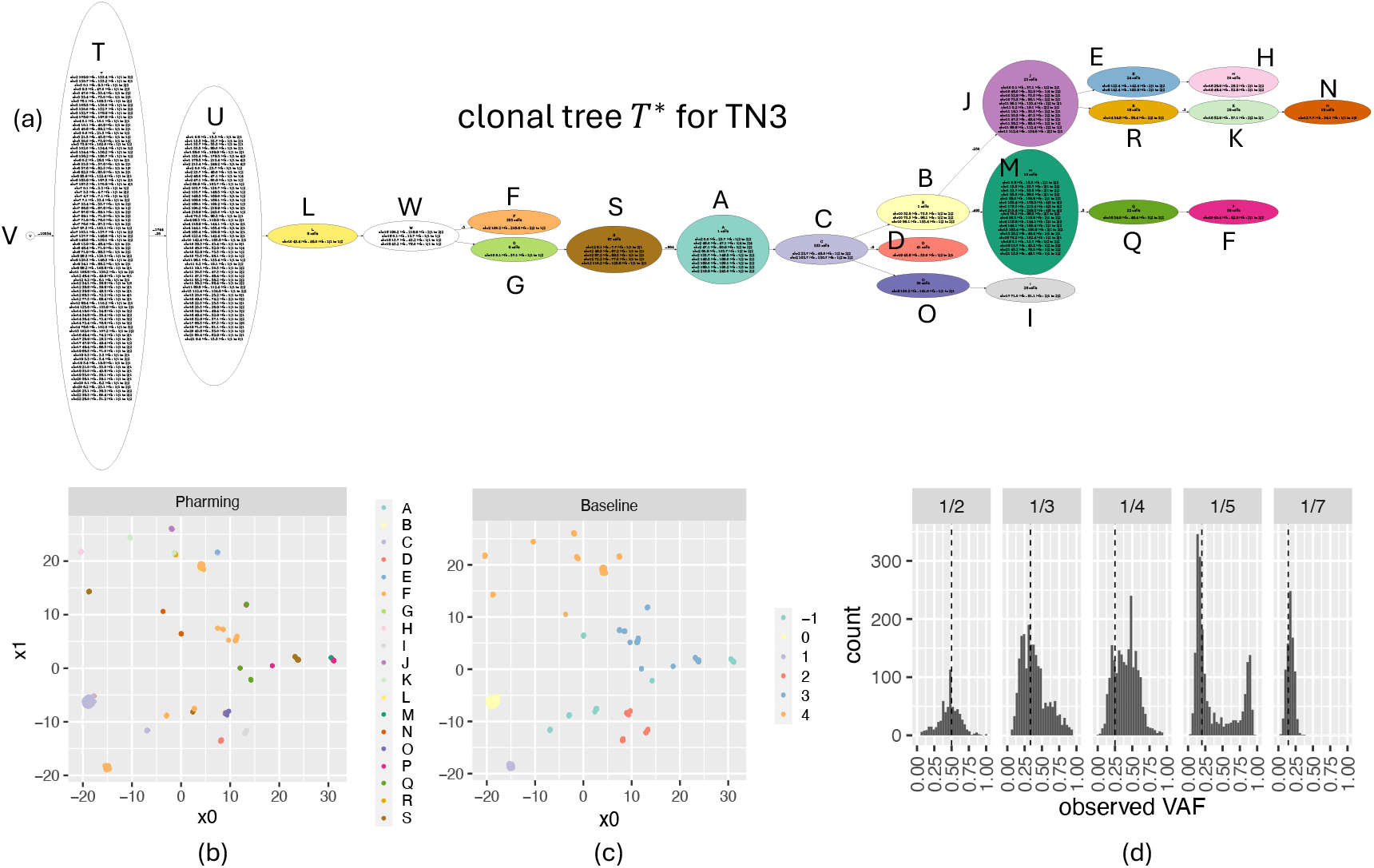
Triple negative breast cancer tumor TN3. (a) Inferred clonal tree *T* with nodes labeled by number of cells and changes in copy number state. (b) UMAP embedding of allele-specific copy number profiles with points colored by Pharming inferred cell mapping *ϕ*. (c) UMAP embedding of allele-specific copy number profiles with points colored by the Baseline density-based clustering. Label −1 indicates a point classified as an outlier. (d) Distribution of observed VAFs for SNVs group by latent VAF given by the genotype.

**Figure S14:**
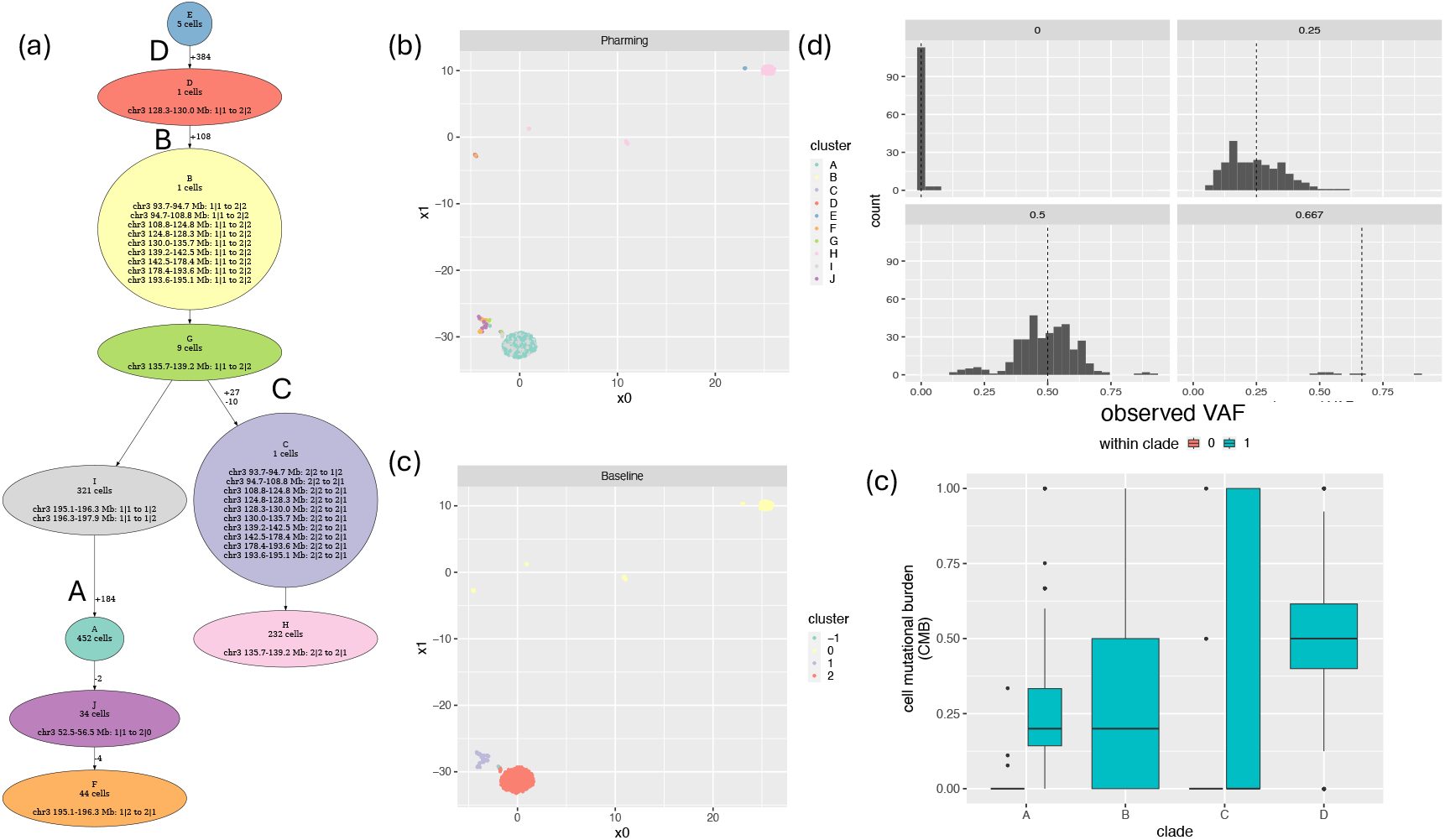
Triple negative breast cancer tumor TN1 clonal tree for chromosome 3. (a) Chromosomal clonal tree *T* ^(3)^ for chromosomal 3 inferred by Pharming with nodes labeled by number of cells and changes in copy number state and edges labeled by the number of SNVs introduced (+) or lost (-). (b) UMAP embedding of allele-specific copy number profiles with points colored by the Pharming inferred cell mapping *ϕ*. (c) UMAP embedding of allelespecific copy number profiles with points colored by the Baseline density-based clustering. Label −1 indicates a point classified as an outlier. (d) Distribution of observed VAFs for SNVs group by latent VAF given by the genotype.(c) Distribution of CMB by clades with newly introduced SNV clusters in the inferred chromosomal clonal trees.

**Figure S15:**
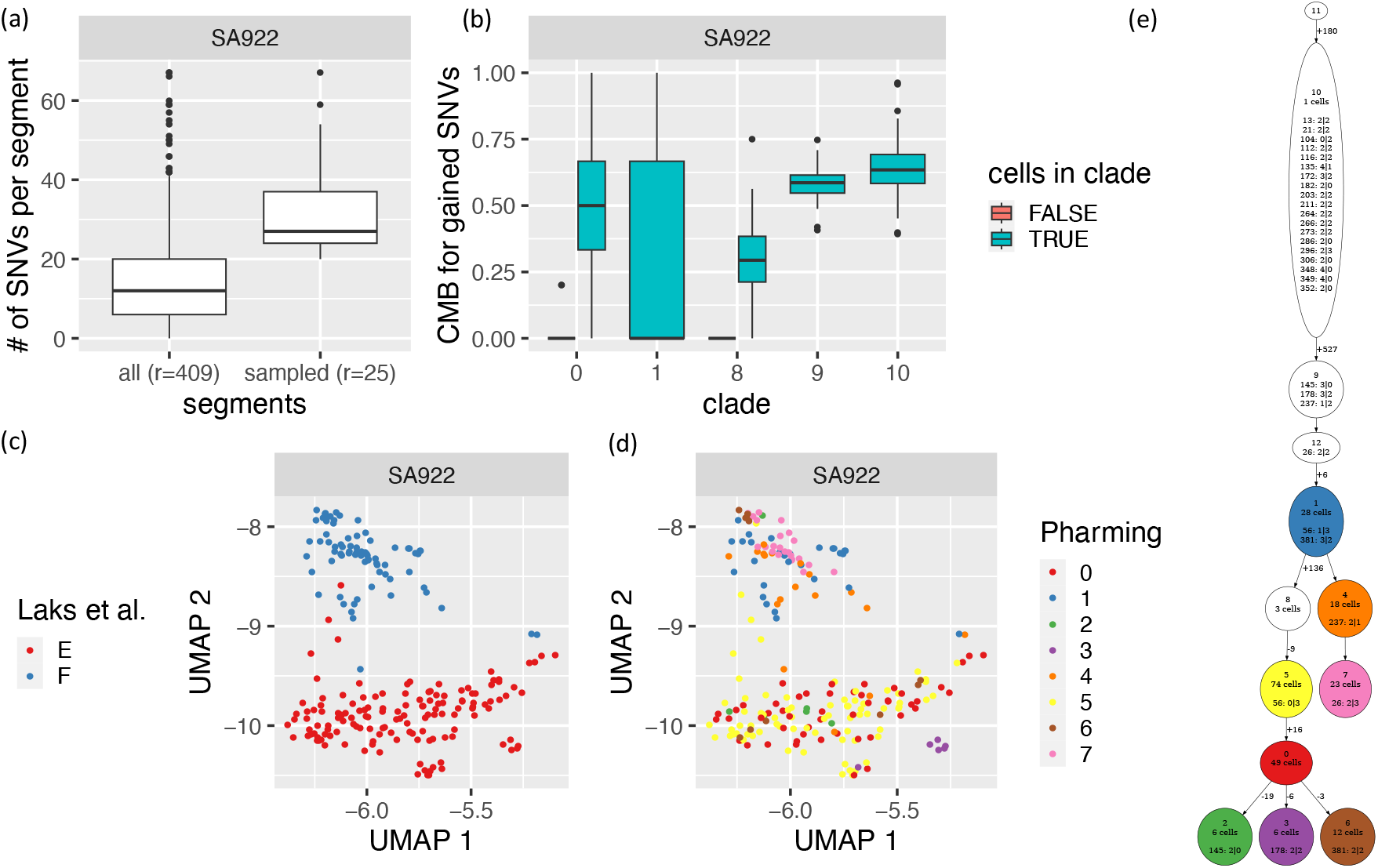
Pharming analysis of the SA922 ovarian cancer cell lines sequenced with DLP+ [13]. (a) Distributions of the number of SNVs in all segments (*r* = 409) and the number of SNVs in the sampled segments (*r* = 25). (b) Distribution of the cell mutational burden (CMB) for the clades in inferred Pharming clonal tree. (c) UMAP projection of the cells in SA922 colored by Laks et al. [13] cluster. (d) UMAP projection of the cells in SA922 colored by Pharming cell assignment *ϕ*. (e) Pharming inferred clonal tree for the 25 sampled segments of SA922. Nodes with more 5 cells assigned are colored to be consistent with (d) and labeled by the number of assigned cells as well as the introduction of a copy number state change, i.e., segment: *x y*. The incoming edges are labeled by the number of newly introduced SNVs and the number of lost of SNVs when applicable.

**Figure S16:**
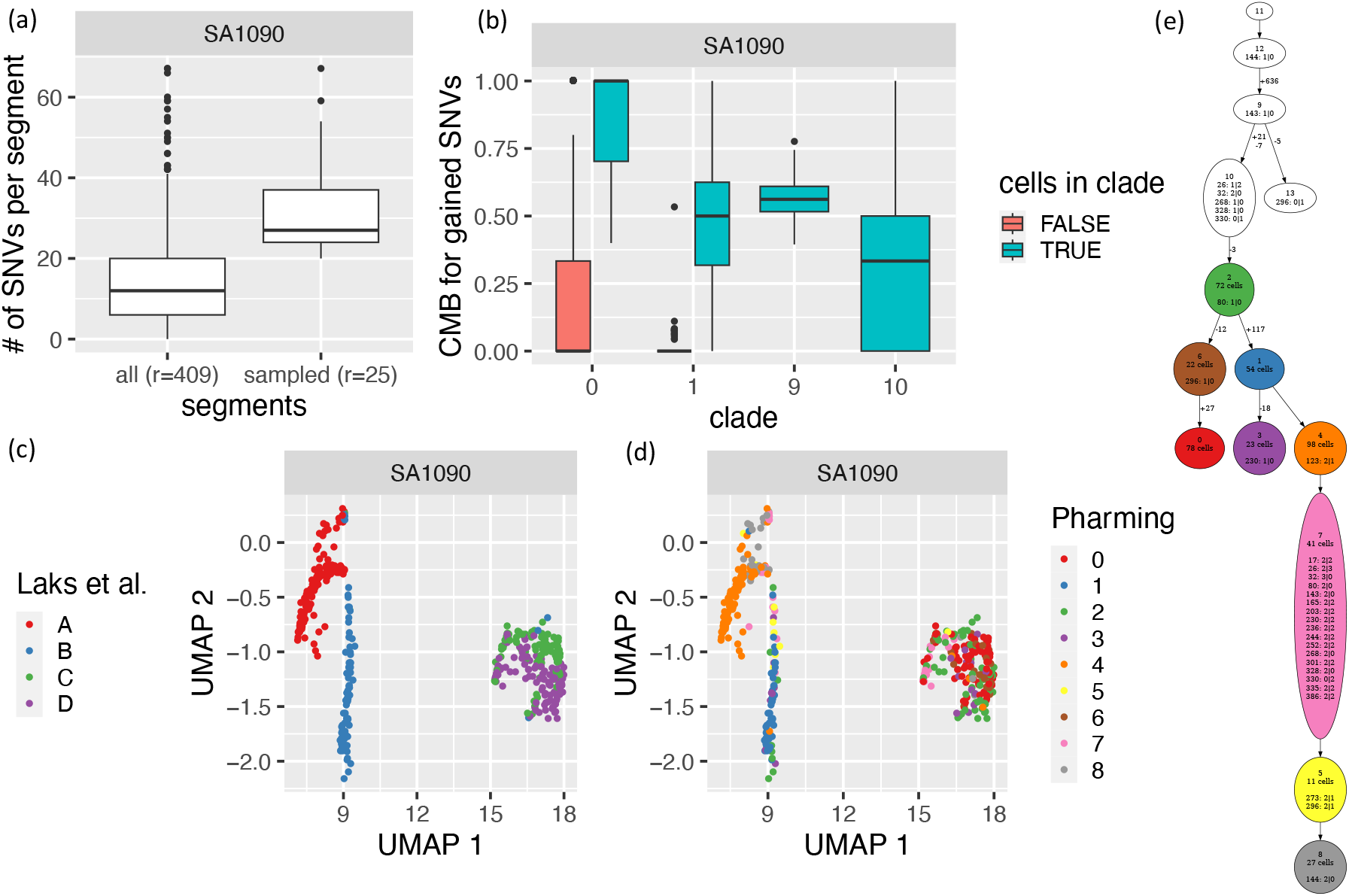
Pharming analysis of the SA1090 ovarian cancer cell lines sequenced with DLP+ [13]. (a) Distributions of the number of SNVs in all segments (*r* = 409) and the number of SNVs in the sampled segments (*r* = 25). (b) Distribution of the cell mutational burden (CMB) for the clades in inferred Pharming clonal tree. (c) UMAP projection of the cells in SA1090 colored by Laks et al. [13] cluster. (d) UMAP projection of the cells in SA1090 colored by Pharming cell assignment *ϕ*. (e) Pharming inferred clonal tree for the 25 sampled segments of SA1090. Nodes with more 5 cells assigned are colored to be consistent with (d) and labeled by the number of assigned cells as well as the introduction of a copy number state change, i.e., segment: *x* |*y*. The incoming edges are labeled by the number of newly introduced SNVs and the number of lost of SNVs when applicable.

**Figure S17:**
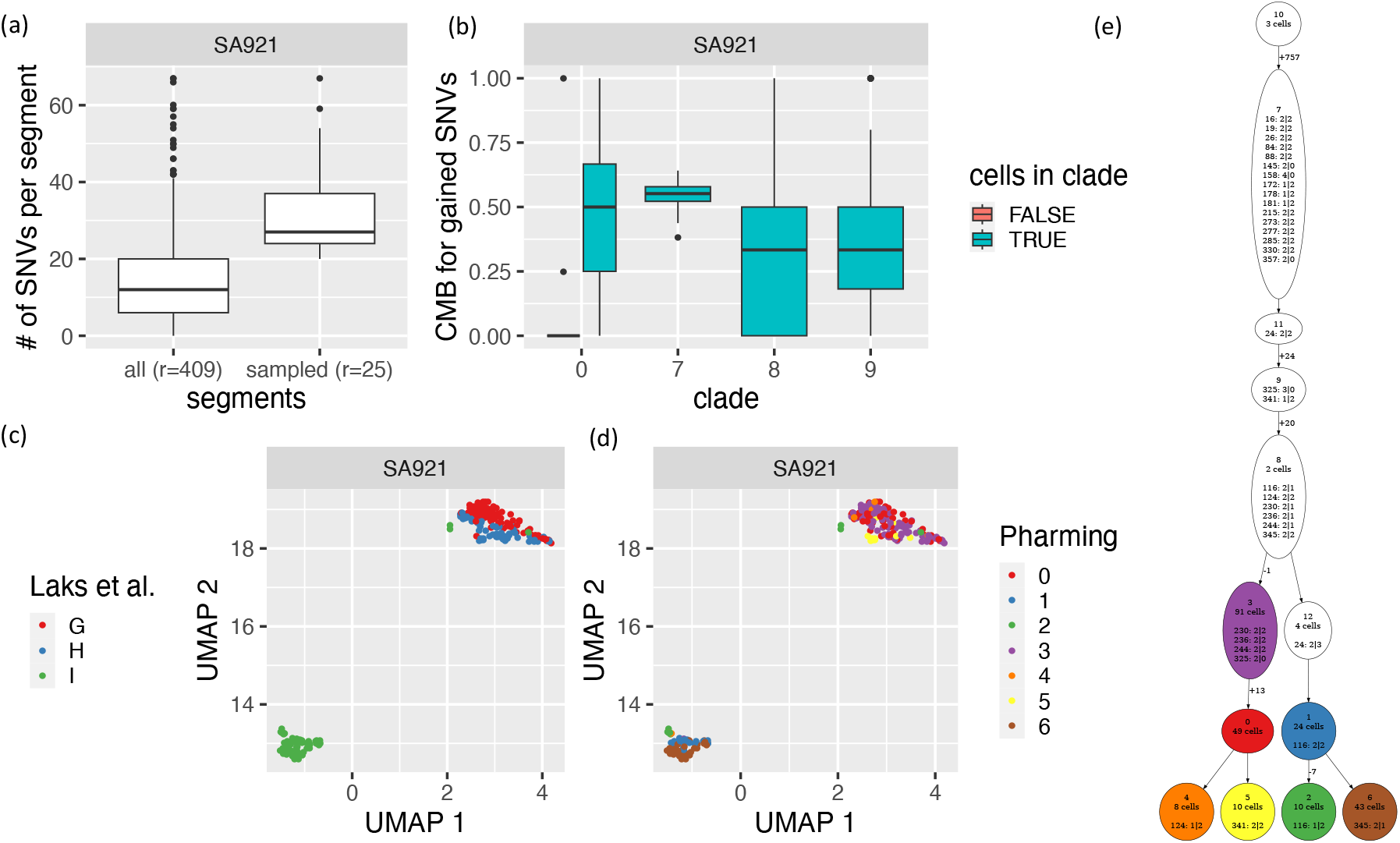
Pharming analysis of the SA921 ovarian cancer cell lines sequenced with DLP+ [13]. (a) Distributions of the number of SNVs in all segments (*r* = 409) and the number of SNVs in the sampled segments (*r* = 25). (b) Distribution of the cell mutational burden (CMB) for the clades in inferred Pharming clonal tree. (c) UMAP projection of the cells in SA921 colored by Laks et al. [13] cluster. (d) UMAP projection of the cells in SA921 colored by Pharming cell assignment *ϕ*. (e) Pharming inferred clonal tree for the 25 sampled segments of SA921. Nodes with more 5 cells assigned are colored to be consistent with (d) and labeled by the number of assigned cells as well as the introduction of a copy number state change, i.e., segment: *x*| *y*. The incoming edges are labeled by the number of newly introduced SNVs and the number of lost of SNVs when applicable.

